# Programmable and Contractile Materials Through Cell Encapsulation in Fibrous Hydrogel Assemblies

**DOI:** 10.1101/2021.04.19.440470

**Authors:** Matthew D. Davidson, Margaret E. Prendergast, Ehsan Ban, Karen L. Xu, Gabriel Mickel, Patricia Mensah, Abhishek Dhand, Paul A. Janmey, Vivek B. Shenoy, Jason A. Burdick

## Abstract

The natural extracellular matrix (ECM) within tissues is physically contracted and remodeled by cells, allowing the collective shaping of functional tissue architectures. Synthetic materials that facilitate self-assembly similar to natural ECM are needed for cell culture, tissue engineering, and *in vitro* models of development and disease. To address this need, we develop fibrous hydrogel assemblies that are stabilized with photocrosslinking and display fiber density dependent strain responsive properties (strain-stiffening, alignment). Encapsulated mesenchymal stromal cells locally contract low fiber density assemblies, resulting in macroscopic volumetric changes with increased cell densities and moduli. Due to properties such as shear-thinning and self-healing, assemblies can be processed into microtissues with aligned ECM deposition or through extrusion bioprinting and photopatterning to fabricate constructs with programmed shape changes due to cell contraction. These materials provide a synthetic approach to mimic features of natural ECM, which can now be processed for applications in biofabrication and tissue engineering.

## Introduction

Natural fibrous matrices, such as fibrin and collagen, are components of the extracellular matrix (ECM) and are widely used *in vitro* for applications such as gel contraction assays in mechanobiology (*1*), self-assembly of vasculature (*2*) and microtissues (*3*), and emerging biofabrication and tissue engineering techniques (*4*–*6*). The use of ECM-derived materials is motivated by their inherent bioactivity, and ability to integrate with, rather than restrict cellular processes (*7*). For example, cell contractility is responsible for many self-assembly or morphogenic processes *in vivo*, such as shaping tissues in embryonic development (*8*) and closing wounds during healing (*9*). Tissue contraction is mediated by the collective action of cells linked together through a web-like network of fibrous ECM molecules, such as collagen and fibrin. Fibrous ECM has structural (e.g., subcellular-scale fiber diameters, micron-scale porosity) (*10*) and mechanical properties (e.g., strain stiffening behavior) (*11*) that support cell contraction and mechanical interactions between cells through the matrix (*12*).

Although useful in many applications, natural matrices have stringent conditions for assembly and may be difficult to control with respect to biochemical and biophysical properties, which limits their reproducibility and utility in many applications (*7*). Further, other ECM derived matrices, such as Matrigel and decellularized ECM are widely used in cell culture, tissue engineering, and even as clinical therapies; however, their batch variability and animal sourcing are limiting (*13*). As an alternative, tunable synthetic materials that closely mimic ECM properties and facilitate self-assembly processes offer a high level of control for the development of tissues *in vitro* and the study of cell-ECM interactions.

Numerous fibrous materials have been engineered to mimic the structure of natural ECM, including self-assembled peptides, electrospun scaffolds, and large-scale fibers. Self-assembled nanofibrous gels recapitulate the structure and scale of ECM fibrils, the smaller precursor to fibers, found in various tissues (*14*). A new class of synthetic amphipathic monomers self-assemble into semiflexible filaments with diameters and persistence lengths similar to cytoskeletal polymers, and form networks with similar strain-stiffening rheology (*15, 16*). These materials can also be coupled with cell adhesion peptides and have been used for cell culture (*17, 18*). Although these gels permit cell encapsulation, their nano-scale porosity limits cell mobility and it is unclear if cells can contract these materials similar to natural ECM (*14*). Micro-to-nano scale fibrous materials that permit cell encapsulation and have diameters within the range of ECM fibers (*14*) have been described; however, the materials used in these systems are highly rigid, providing fibers that cells cannot physically remodel (*19, 20*). Another important material is formed by electrospinning hydrogel fibers with moduli similar to natural ECM that allow cells to contract and compact fibers in a manner similar to collagen contraction; (*21*) yet, these electrospun scaffolds are rather stiff pre-formed solids that are not amenable to cell encapsulation or injection. Lastly, supracellular scale synthetic fibrous (*22*) or ribbon-like (*23*) materials that permit cell encapsulation and approximate the size of large collagen fibers (*14*) have also been described; however, their large sizes hinder multi-cellular interactions through the matrix. Thus, synthetic systems with improved control over structure and mechanics would provide the field with tools to study cell-ECM interactions and engineer self-assembled tissues through combined top-down and bottom-up approaches.

To overcome the limitations of current engineered systems, we report here on fibrous hydrogel assemblies composed of micro-to-nano diameter fibers that are fabricated from diverse materials, are injectable, and exhibit strain stiffening behavior based on fiber density. Additionally, this system permits cell encapsulation and contractility, features that allow assembly into microtissues and the fabrication of cell-laden constructs. These materials are programmable by altering the extent of contractility through fiber density, the shape of constructs through patterning, and dynamic shape changes with culture through the fabrication of multi-material constructs.

## Results

### Generation of photocrosslinkable fibrous hydrogel assemblies

To recreate the 3D architecture of fibrous ECM, we introduce a method to fabricate suspensions of hydrogel fibers that can be assembled and crosslinked into a highly porous fibrous network (Fig 1a). This process involves: (1) mechanical fragmentation of electrospun hydrogel fiber mats into fiber suspensions, (2) assembly of fibers into networks through fiber concentration at desired densities, and (3) stabilization of fiber networks through photocrosslinking to form fibrous hydrogel assemblies.

**Figure 1.**
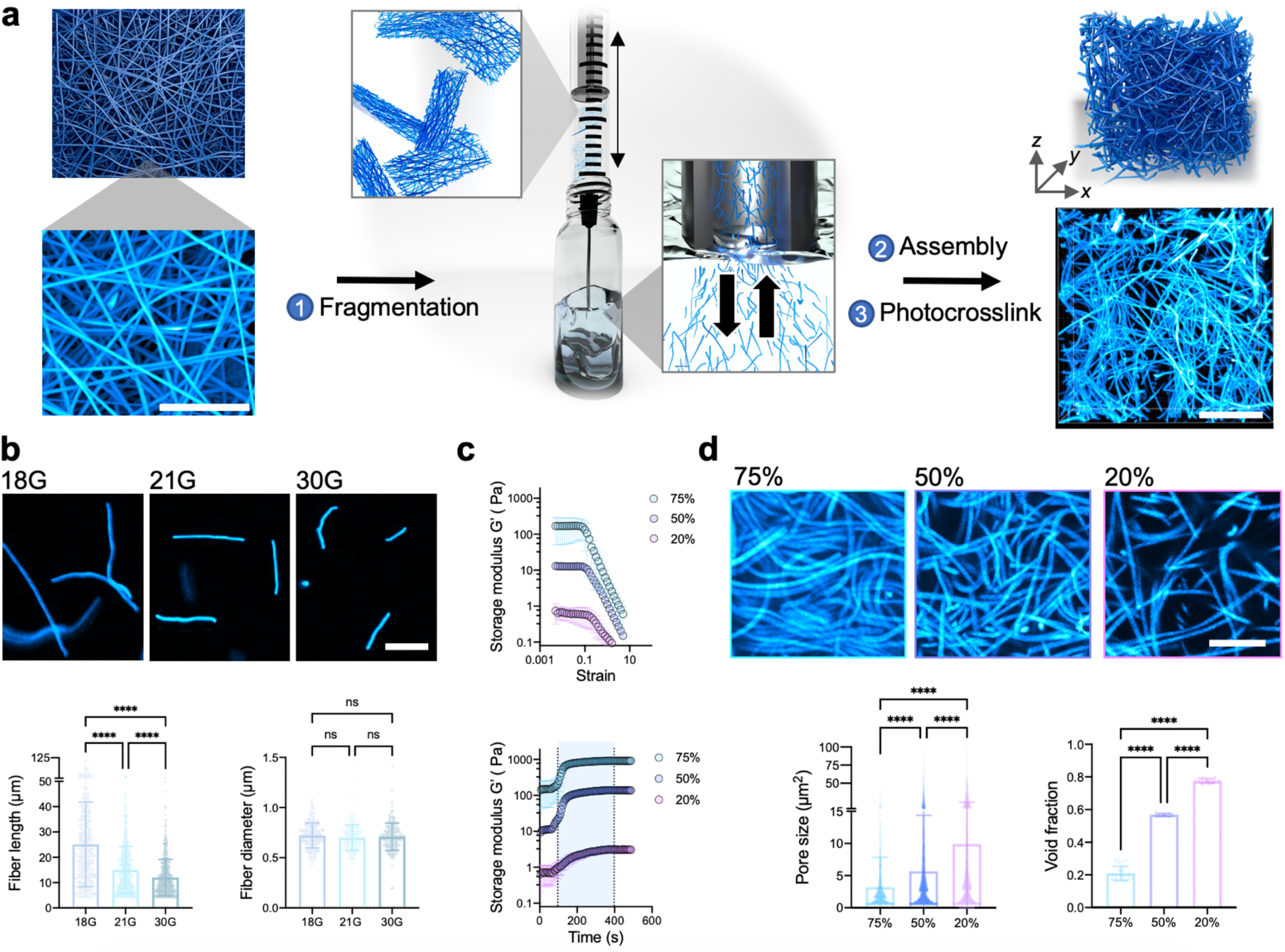
Fabrication of fibrous hydrogel assemblies. **(a)** Schematics (left top, center, and right top), as well as scanning electron microscopy (left bottom) and fluorescent (right bottom) images demonstrating: (1) the production of fiber suspensions from electrospun scaffolds via fragmentation by passing scaffold pieces repeatedly through a syringe needle, (2) assembly of fibers into a fibrous network, and (3) photocrosslinking of the fibrous network for stabilization through inter-fiber crosslinking. Images are representative of n=3 independent samples. Scalebars 5 µm (left), 25 µm (right). **(b)** Representative fluorescent images (top) and quantification of fiber length (bottom left) and diameter (bottom right) for fiber suspensions obtained from fragmentation with 18G, 21G, or 30G needles. Images and data are representative of n=3 independent experiments (fiber length, mean ± s.d., one-way ANOVA, 18G vs. 21G *p* = 1.0 x 10^−15^, 18G vs. 30G *p* = 1.0 x 10^−15^, 21G vs. 30G *p* = 5.0 x 10^−7^). Scalebar 25 µm. **(c)** Storage modulus (G’) of assemblies before photocrosslinking with increasing strain (top) and during photocrosslinking (bottom, blue region denotes visible light exposure at 5 mW/cm^2^) at varied fiber densities (75%, 50%, 20%) (mean ± s.d., n=3 independent samples). **(d)** Representative fluorescent images (top) and quantification of pore size (µm^2^) (bottom left) and void fraction (bottom right) of assemblies at varied fiber densities (75%, 50%, 20%). Images and data are representative of n=3 independent samples ((pore size, mean ± s.d., one-way ANOVA, 18G vs. 21G *p* = 1.0 x 10^−7^, 18G vs. 30G *p* = 1.0 x 10^−7^, 21G vs. 30G *p* = 5.0 x 10^−7^), (void fraction, mean ± s.d., one-way ANOVA, 18G vs. 21G *p* = 2.0 x 10^−11^, 18G vs. 30G *p* = 2.0 x 10^−11^, 21G vs. 30G *p* = 2.0 x 10^−11^)). Scalebar 25 µm. (n.s. not significant, ****p<0.0001).

The fragmentation process is a simple approach to fabricate fiber suspensions from electrospun scaffolds, primarily in this study with norbornene modified hyaluronic acid (NorHA) (Supplementary Fig 1). NorHA has been routinely processed into fibrous hydrogel scaffolds through electrospinning, using thiol-ene reactions to stabilize scaffolds with both inter-fiber and intra-fiber crosslinking (*24, 25*). Electrospun scaffolds are limited in that they are not injectable and cannot be used to encapsulate cells; however, fragmentation converts these scaffolds into hydrogel fiber suspensions that overcome these limitations. Specifically, electrospun scaffolds are cut into small pieces, briefly hydrated, passed repeatedly through a syringe needle, and purified from any larger fragments with filtration, which disrupts inter-fiber crosslinks and fractures fibers along their length (Supplementary Fig 2). There are many parameters that can be varied in the fragmentation process and electrospun scaffold properties to influence final fiber lengths, such as the needle gauge used to fragment fibers or the extent of crosslinking of the fibers. We observed that increased needle gauge or reduced crosslinking results in fibers with decreasing lengths, while maintaining fiber diameters (Fig 1b, Supplementary Fig 3). Further, the fragmentation process is compatible with electrospun scaffolds from a wide range of polymers (e.g., gelatin (NorGel), poly(ethylene glycol) (NorPEG), HA (MeHA)), which illustrates the versatility of the technique (Supplementary Fig 4).

To improve upon fiber yield, we implement a multi-fiber electrospinning process, where sacrificial fibers of a dissolvable poly(ethylene oxide) are electrospun with stable NorHA fibers (Supplementary Fig 5). The sacrificial fiber population physically blocks inter-fiber bonds during scaffold fabrication, but then dissolves away, allowing for easier fragmentation (*26*). Overall, this process results in reproducible NorHA fibers with lengths similar to those reported for natural ECM (*27*) (∼25 µm) and ECM-like fiber diameters (*14*) (∼700 nm) (Fig 1b, Supplementary Fig 5). For the remaining studies, NorHA fibers (∼50% consumption of norbornene groups based on calculated level of norbornene modification) fabricated using the multi-fiber electrospinning approach and fragmentation (18G, then 21G needle, with 40 passes through each needle) are used. This approach provides a rapid and efficient method to generate fiber suspensions and overcomes challenges with alternate approaches, such as issues with light diffusion and polymer loss using photopatterning (*28*), lengthy multi-step protocols for cryosectioning (*29*), or harsh chemical treatments with aminolysis (*30*).

After production, fiber suspensions are centrifuged and resuspended at desired concentrations (75%, 50%, 20% v/v) to form fiber assemblies. These assemblies behave as yield stress fluids at all fiber densities investigated, showing yielding behavior with increased strain (Fig 1c). Additionally, the assemblies are injectable and exhibit shear-thinning and self-healing behavior, represented as a reduction in viscosity with increasing shear rate and a loss and recovery of storage moduli across cycles of low and high strains, likely due to fiber entanglement and packing during processing (Supplementary Fig 6). After assembly, remaining norbornene groups on the fibers can be used for an additional thiol-ene reaction to stabilize the networks. Specifically, additional photoinitiator (lithium phenyl-2,4,6-trimethylbenzoylphosphinate (LAP)) and crosslinker (4-arm PEG-thiol, 5kDa) are added to the assemblies, which then photocrosslink with exposure to visible light (400-500 nm λ), measured as an increase in storage modulus (G’) (75%: 146.4 to 915.6 Pa, 50%: 10.5 Pa to 138.2 Pa, 20%: 0.7 Pa to 3.1Pa) (Fig 1c). Importantly, fiber densities below 20% are unstable after crosslinking, with bulk gel dissolution during hydration. Although other chemistries could be used to introduce inter-fiber bonds, such as physical crosslinks or dynamic covalent chemistry (*25*), the norbornene chemistry and thiol-ene reaction results in stable networks. These fibrous hydrogel assemblies, referred to as fibrous assemblies from here onward, have pore sizes within the range of fibrous ECM (*14, 31*) (∼3-10 µm^2^) and void fractions ranging from ∼21-78% based on initial fiber volume percentage (Fig 1d).

### Strain responsive properties and modeling of fibrous hydrogel assemblies

Cells physically strain and compact fibrous ECM during tissue contraction, motivating the introduction of strain responsive properties into engineered fibrous assemblies. Importantly, the micron-scale pore size of ECM is thought to contribute to their force responsive properties, where strain causes fibers to stretch and buckle in the direction where stresses are compressive, leading to fiber alignment in the direction of strain and non-linear stiffening (*10, 11*). Using rheological shear strain sweeps, we observe that fibrous assemblies exhibit non-linear mechanical properties, undergoing strain-induced increases in G’ (i.e., strain-stiffening) after reaching a critical strain (γ_c_) (Fig 2a). This is particularly pronounced with the lowest fiber density formulation of 20% volume fraction, which results in the highest relative increase in stiffness (2.3 fold) before failure, whereas the 50% and 75% volume fractions exhibit a lower increase in stiffness (∼1.6 and 1.3 fold, respectively) (Fig 2a). The critical strains (γ_c_) of fibrous assemblies (∼4-6% strain) are similar to those measured with natural fibrous ECM and are within the range of strains that cells impose on the ECM (maximum of ∼30-50% strain), suggesting that cellular strains could potentially stiffen these materials (*32*).

**Figure 2.**
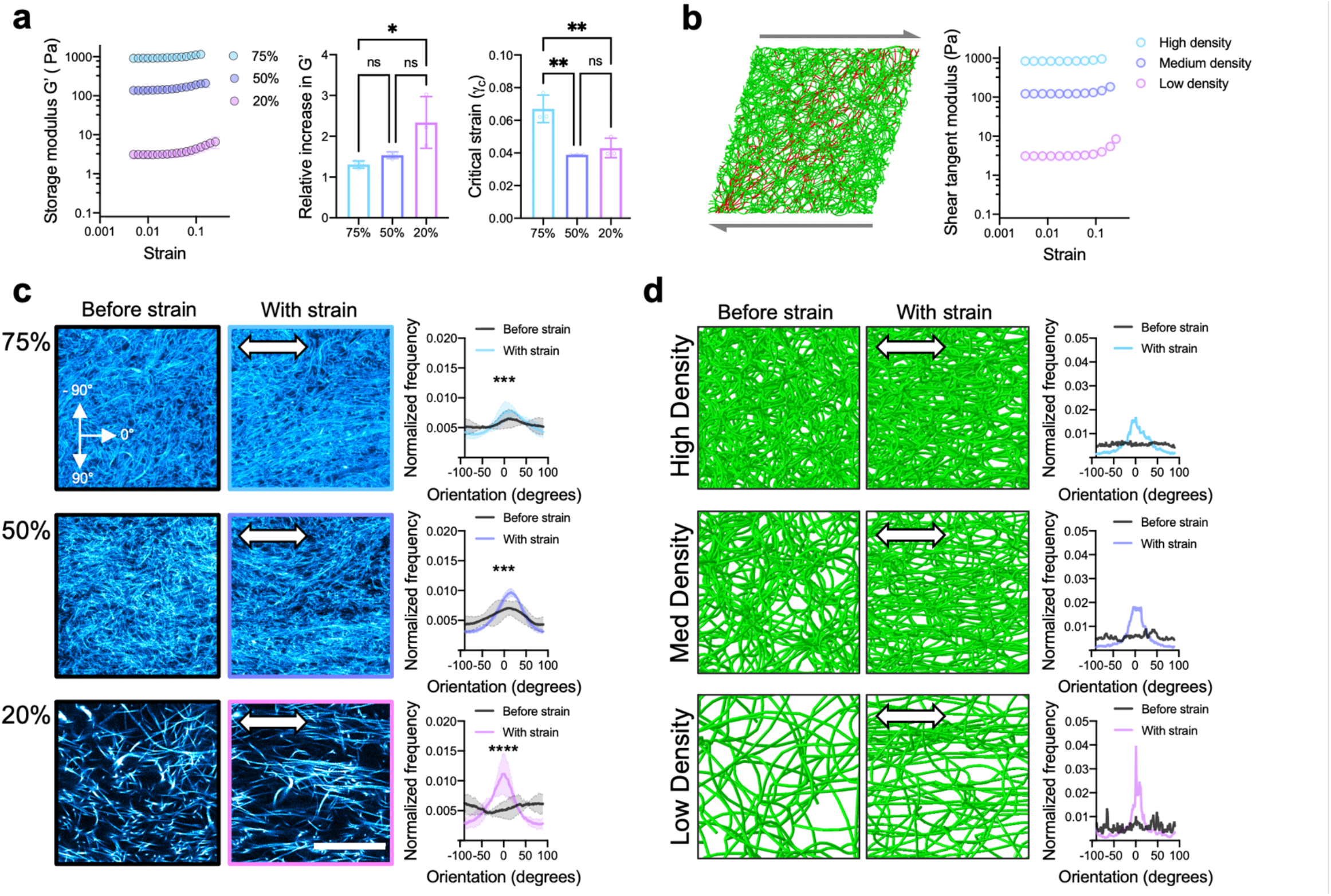
Strain responsive behavior of fibrous hydrogel assemblies. **(a)** Storage modulus (G’) with increasing strain (left, 0.05-0.5 strain, 1 Hz), relative increase in G’ before failure (center, normalized to modulus at low strain, 0.05), and critical strain (γ_c_, onset of stiffening) of assemblies at varied fiber densities (75%, 50%, 20%) (n=3 gel samples, mean ± s.d., one-way ANOVA, (Relative increase in G’, 75% vs. 20% *p* = 0.034), (γ_c_, 75% vs. 50% *p* = 0.003, 75% vs. 20% *p* = 0.006)). **(b)** Snapshot of discrete fiber network model deforming under shear strain (left, red fibers indicate stretched fibers) and model tangent modulus with increased shear strain (right, 0.05-0.5 strain) at varied network densities (High, Medium, Low). **(c)** Representative fluorescent images (before (left column) and with uniaxial strain (center column, ∼0.1 strain)) and quantification of fiber orientation (right column) within assemblies before and with strain at varied fiber densities (75%, 50%, 20%) (n=3 samples, mean ± s.d., Watson-Wheeler test for homogeneity, 75% *p* = 0.0005, 50% *p* = 0.0005, 20% *p* = 7.0 x 10^−6^). Scalebar 25 µm. **(d)** Representative snapshots of fiber network model before (left column) and with simulated uniaxial strain (center column) and profiles of fiber orientation (right column) in model networks before and with strain at varied fiber densities (High, Medium, Low). (n.s. not significant, *p<0.05, **p<0.01, ***p<0.001, ****p<0.0001).

To further understand the influence of fiber density on strain stiffening, we introduce a discrete fiber network computational model based on the structure and mechanics of fibrous assemblies and imposed shear strains on the networks (Fig 2b). The fiber network model shows similar trends to our experimental data, with low density fiber networks exhibiting higher relative strain-stiffening when compared to high-density fiber networks. During shear tests with the fiber network model, we observe higher levels of fiber bending and reorientation with low density networks, suggesting that fibers reorient within fibrous assemblies during strain. To investigate how imposed strains change fibrous assembly micro-architecture, we use confocal microscopy to image fiber network architectures before and during uniaxial strain (10%) (Fig 2c, Supplementary Fig 7). Fibrous assemblies undergo strain-induced alignment, with low fiber densities (20%) having the highest relative increase in alignment, whereas higher fiber densities (50%, 75%) had similar levels of alignment relative to networks that are not strained (Fig 2c). Importantly, when uniaxial strains (40% strain) are imposed on the discrete fiber network model, high levels of fiber re-alignment are observed in the direction of strain with the greatest alignment observed for low density networks (Fig 2d).

Our engineered fiber assemblies are unique in their ability to mimic feature sizes and architecture of reconstituted fibrous ECM, while also introducing parameters of injectability and tunable non-linear properties. While other non-linear synthetic material systems have been described (*15, 33*–*35*), these materials fail to capture the micron-scale porosities, fiber diameters, lengths, and 3D architectures that we observe with these fibrous hydrogel assemblies. Importantly, these results are successfully modelled to describe our experimental observations of strain stiffening and fiber alignment with variations in fiber concentration. Our results are also similar to what has been reported for type I collagen gels produced with low and high collagen concentrations (*36, 37*), are consistent with previous computational models (*12, 31*), and further validate the ECM-like properties of our fibrous assemblies.

### Cell-mediated fiber recruitment and contractility of fibrous hydrogel assemblies

Having demonstrated that fibrous assemblies adapt to forces similar to native ECM, we next explore the interface between cells and fibers when encapsulated within fibrous assemblies, including features such as cell-mediated fiber contraction and recruitment. A major advantage of these fibrous assemblies over existing electrospun hydrogel scaffolds is that cells can be encapsulated within the assemblies by mixing with fiber suspensions at desired fiber densities and then photocrosslinking (Fig 3, Supplementary Fig.8). Alternative fiber systems that allow for cell encapsulation exist, but they employ rigid polymers that cells cannot remodel (*19, 20*), are composites with extensible fibers embedded in a continuous gel phase that limits fiber recruitment (*28*), or are on scales that are much larger than single cells (*22, 23*). To engage cell adhesion through integrins, fibrous assemblies are decorated with a peptide containing the adhesive RGD motif for all cell studies. Human mesenchymal stromal cells (MSCs), a prototypical cell type used in mechanobiological studies and of great interest in tissue engineering, are encapsulated within fibrous assemblies of 3 different fiber densities (75%, 50%, and 20%), and fiber recruitment and cell morphology are assessed during culture (Fig 3 a-c). MSCs spread and recruit fibers to the cell surface dynamically within 3 days in culture in low fiber densities (20%), whereas fiber recruitment is less apparent in higher fiber densities (Fig 3b). At high fiber densities, there are likely impediments to fiber recruitment due to the high fiber concentrations and resulting greater mechanical resistance. Interestingly, there are no statistically significant differences in cell aspect ratio and circularity across the 3 fiber volume fractions after 3 days in culture, suggesting that cells are able to spread within all fibrous assemblies, regardless of initial conditions.

While cell morphology is similar across the different initial fiber densities, we observe large differences in the levels of fiber recruitment. This is quantified by measuring the average fiber fluorescence intensity in 2D radial bands around cells (using 1 µm increments) to define a fiber compaction length that can be compared to remote fiber locations (Fig 3d, Supplementary Fig 8). Cells in low fiber densities (20%) compact large fiber “shells” (∼21 µm in radius) around the cell surface, while fiber compaction length decreases with medium and high fiber densities (∼12µm, and ∼2.5 µm shell radius for 50% and 75%, respectively) (Fig 3e,f). These results are supported by the previous experiments and simulations with our fiber network model, which show that the smaller pores and higher elastic moduli in high density fiber formulations limit the extent of fiber reorganization that can occur, while large pores support fiber reorganization and compaction. These results are also similar to what has been observed for cells cultured atop 2D fibrous networks (*21*), but we now consider the more challenging cultures of cells in 3D that is often more physiologically relevant.

**Figure 3.**
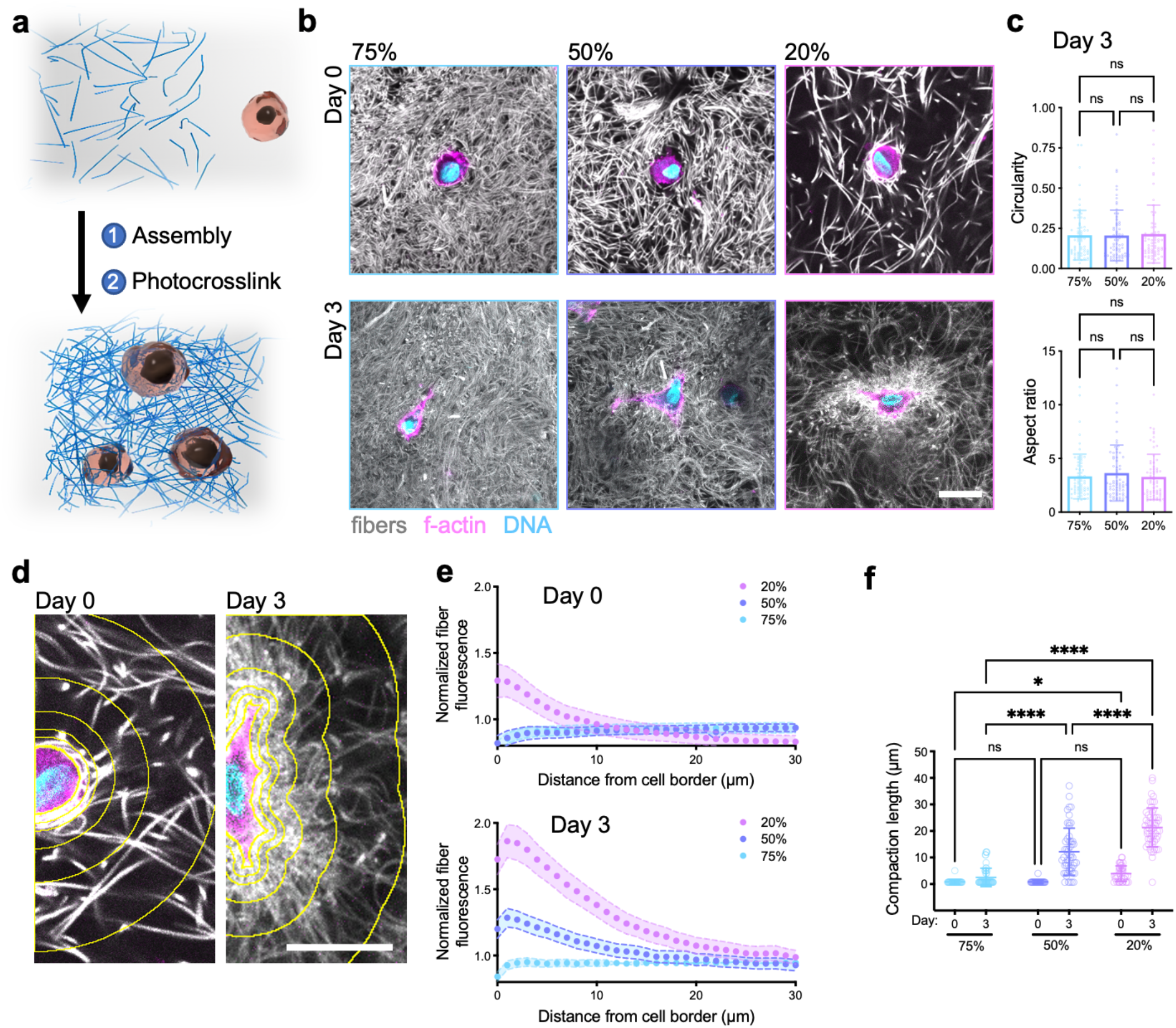
Mesenchymal stromal cell behavior within fibrous hydrogel assemblies. **(a)** Schematic of cell encapsulation process through fiber assembly and photocrosslinking and **(b)** representative fluorescence images of encapsulated cells (actin (pink), nuclei (blue), fibers (grey)) at day 0 (top) and day 3 (bottom) of culture at varied fiber densities (75%, 50%, 20%). Images are representative of n=3 biologically independent samples. **(c)** Quantification of cell morphology (circularity, aspect ratio) from max z-projections of encapsulated cells at day 3 of culture (n=67, 69, 67 cells from 3 biologically independent experiments, mean ± s.d., one-way ANOVA). **(d)** Representative images (shown for 20% densities) with progressive bands around cells used to quantify fiber fluorescence intensity with distance from the cell-gel interface and **(e)** profiles of normalized fiber fluorescence at days 0 (top) and 3 (bottom) of culture (normalized to fluorescence intensity in a remote area without cells) at varied fiber densities (75%, 50%, 20%) (Day 0: n=34, 28, 45 cells from 3 biologically independent experiments for 75%, 50% and 20% respectively, mean ± s.d.; Day 3: n=53, 54, 52 cells from 3 biologically independent experiments for 75%, 50% and 20% respectively, mean ± s.d.). **(f)** Quantification of fiber compaction length (distance away from cell border where compacted fiber fluorescence intensity reaches background intensity level) at varied fiber densities (75%, 50%, 20%) on day 0 and day 3 of culture (Day 0: n=34, 29, 38 cells from 3 biologically independent experiments for 75%, 50% and 20%, respectively; Day 3: n=53 cells from 3 biologically independent experiments; mean ± s.d., mixed model analysis; Day 0: 75% vs. 20% *p* = 0.048; Day 3: 75% vs. 50% *p* = 1.5 x 10^−13^, 75% vs. 20% *p* = 1.2 x 10^−13^, 50% vs. 20% *p* = 1.6 x 10^−13^). Scalebars 25 µm. (n.s. not significant, * p<0.05, ****p<0.0001).

Having established that cells can locally compact fibrous assemblies, we next explore if the collective contraction of cells at higher densities results in macroscale contraction (Fig 4a). When MSCs are encapsulated at high densities (1×10^7^ cells ml^-1^) and the gel area monitored over time, we observe high cell viability and bulk contraction of constructs fabricated with low fiber densities (20%) within 1 day of culture (∼36% of original area), while contractions are greatly reduced with increasing fiber densities (∼85 and 95% of original area for 50 and 75% fiber densities, respectively) (Fig 4b-d, Supplementary Fig 9). After 3 days of culture, contraction stabilizes, and low fiber densities (20%) decrease to ∼12% of their original volume, while higher fiber densities only decrease in volume to ∼51% and 78% of their original volume, for medium (50%) and high (75%) fiber densities, respectively (Fig 4c). As gel volume decreases, the cell density simultaneously increases over seven-fold to 7.1×10^7^ cells ml^-1^ for low fiber densities (20%), whereas the cell density only slightly increases to 1.8×10^7^ and 1.5×10^7^ cells ml^-1^, for medium (50%) and high (75%) fiber densities, respectively (Fig 4e). Additionally, even higher cell densities are obtained after contraction (11.5×10^7^ cells ml^-1^) by initially encapsulating cells at 5×10^7^ cells ml^-1^ within low fiber density (20%) fibrous assemblies (Supplementary Fig 10). These high cell densities may be particularly useful in applications of the engineering of tissues and models for cell-dense tissues.

**Figure 4.**
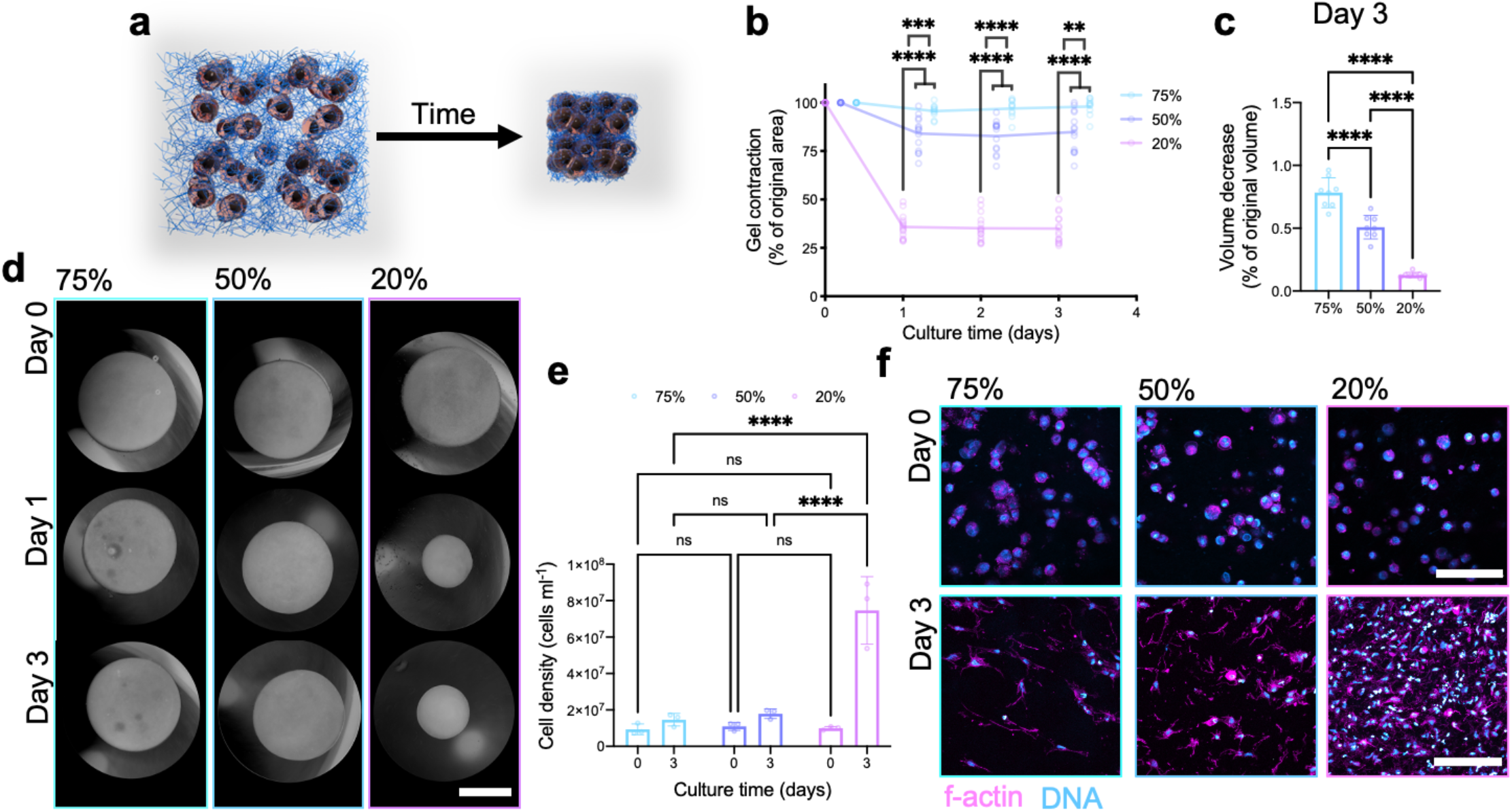
Multicellular contraction of fibrous hydrogel assemblies. (a) Schematic and (b) quantification of assembly contraction (% of original sample area) for MSCs (1 x 10^7^ cells/ml) encapsulated within assemblies with varied fiber densities (75%, 50%, 20%) (n=10, 15, and 15 samples from 3, 4, and 4 biologically independent experiments for 75%, 50% and 20%, respectively, mean ± s.d., two-way ANOVA (Day 1: 75% vs. 50% *p* = 8.0 x 10^−4^, 75% vs. 20% *p* = 4.0 x 10^−15^, 50% vs. 20% *p* = 5.0 x 10^−14^; Day 2: 75% vs. 50% *p* = 7.0 x 10^−5^, 75% vs. 20% *p* = 1.0 x 10^−14^, 50% vs. 20% *p* = 5 x 10^−14^; Day 3: 75% vs. 50% *p* = 0.0014, 75% vs. 20% *p* = 5.0 x 10^−15^, 50% vs. 20% *p* = 8.0 x 10^−13^)). (c) Quantified contraction induced volume decrease at day 3 (relative to day 0 volume) for varied fiber densities (75%, 50%, 20%) (n=8 samples from 2 biologically independent experiments, mean ± s.d., one-way ANOVA, 75% vs. 50% *p* = 1.1 x 10^−5^, 75% vs. 20% *p* = 4.2 x 10^−12^, 50% vs. 20% *p* = 7.5 x 10^−8^). (d) Representative inverted phase contrast images of assemblies over time with varied fiber densities (75%, 50%, 20%). Images are representative of n=3 for 75% and n=4 for 50% and 20% independent experiments. Scalebar 2 mm. (e) Quantification of cell density and (f) representative fluorescent images (actin (pink), DNA (blue)) for MSCs encapsulated within assemblies after 0 or 3 days of culture with varied fiber densities (75%, 50%, 20%) (n=3 biologically independent experiments, mean ± s.d., two-way ANOVA (Day 3: 75% vs. 20% *p* = 2.3 x 10^−6^, 50% vs. 20% *p* = 4.1 x 10^−6^). Images are representative of n=3 biologically independent experiments. Scalebar 100 µm. (n.s. not significant, ** p<0.01, *** p<0.001, ****p<0.0001).

With regards to mechanical properties of fibrous assemblies with contraction, we observe that contracted constructs formed with low fiber densities (20%) have significantly higher compressive moduli (3.3 kPa) after 3 days when compared to constructs formed with higher fiber densities (1.4 and 1.8 kPa, for 50 and 75%, respectively) (Supplemental Fig 11a,b). These results suggest that cells substantially stiffen assemblies with culture when compared to estimated non-contracted fibrous assembly moduli from rheological time sweeps as 0.006 kPa, 0.276 kPa, and 1.826 kPa for 20%, 50%, and 75% fibrous assemblies, respectively. ECM deposition also likely contributes to increased mechanics, as fibronectin is extensively deposited throughout contracted low fiber density constructs (20%), whereas fibronectin is limited to pericellular regions in higher fiber densities (Supplementary Fig 11c).

To further investigate cellular contributions to assembly stiffening with culture, we inhibit cell contraction with blebbistatin (myosin II inhibitor) or remove/lyse cells with SDS and observe that both of these perturbations result in decreased moduli, ∼72% and 49% of non-treated gels, respectively (Supplementary Fig 11d). Plastic deformation of the fibers can also influence mechanics (*38*), and low cell density experiments with blebbistatin and SDS treatment do not result in recoiling of the fibers away from cells, suggesting that cells plastically deform fibrous assemblies (Supplementary Fig 11e). These results further support the ECM-like properties of fibrous assemblies, showing that cells can stiffen assemblies and plastically deform them through contraction (*39, 40*).

### Microtissues via contractile fibrous hydrogel assemblies

Microtissues formed from the cellular embedding and remodeling of ECM have been used to study cell and ECM organization during wound healing (*41*), model healthy and diseased tissue (*42, 43*), and promote iPSC maturation (*44*). This bottom-up approach relies on stromal cell ECM contraction and physical interactions with the matrix to form high cell density tissue mimics (*3, 45*). Although we have learned a great deal from current microtissue systems, there is a need for systems where the matrix can be tuned to better understand the contribution of ECM properties to tissue morphogenesis (*45*) and ECM-based diseases such as fibrosis (*46*). Currently, most commercially available ECM systems (e.g., type I collagen, fibrin, Matrigel) have limited control over biochemical and biophysical properties and would benefit from synthetic versions where such properties can be modulated.

To address this need, we assess the potential for the fabrication of microtissues using fibrous assemblies, where cell contraction results in the formation of aligned tissues in the direction of principal stress dictated by micropost geometry (Fig 5a). Micropost arrays with 1-, 2-, and 3-post geometries are fabricated to give tissues with cylindrical, uniaxial, and random alignment, respectively. Having observed extensive contraction with low fiber densities (20%), we use these formulations for microtissue formation. Microtissues form over 24 hours, resulting in circular, dog bone, and triangular-shaped tissues that can be cultured for 14 days to allow for collagen I deposition with alignment that corresponds to macroscale tissue geometry (Fig 5b,c, Supplementary Fig 12). Specifically, 1-post geometries result in microtissues with circumferentially aligned collagen, 2-post geometries result in highly aligned collagen throughout the center of the microtissue, and 3-post geometries have randomly aligned collagen in the center of microtissues. ECM alignment dictates cell shape and function, such as with circumferential alignment within arterial walls supporting dynamic changes in volumes, load bearing tissues such as tendons needing highly aligned ECM, and glandular tissues such as the liver require isotropic ECM to support filtration. By introducing mechanical constraints, cells embedded within fiber assemblies transition from an initial isotropic orientation into organized tissues that mimic the ECM architecture found in some tissues. Importantly, the 3D fibrous networks remain within the tissue after 14 days of culture, suggesting that these fibers not only facilitate but also integrate with the microtissue, which could be useful to provide further control over contracted microtissue differentiation and maturation (*47*).

**Figure 5.**
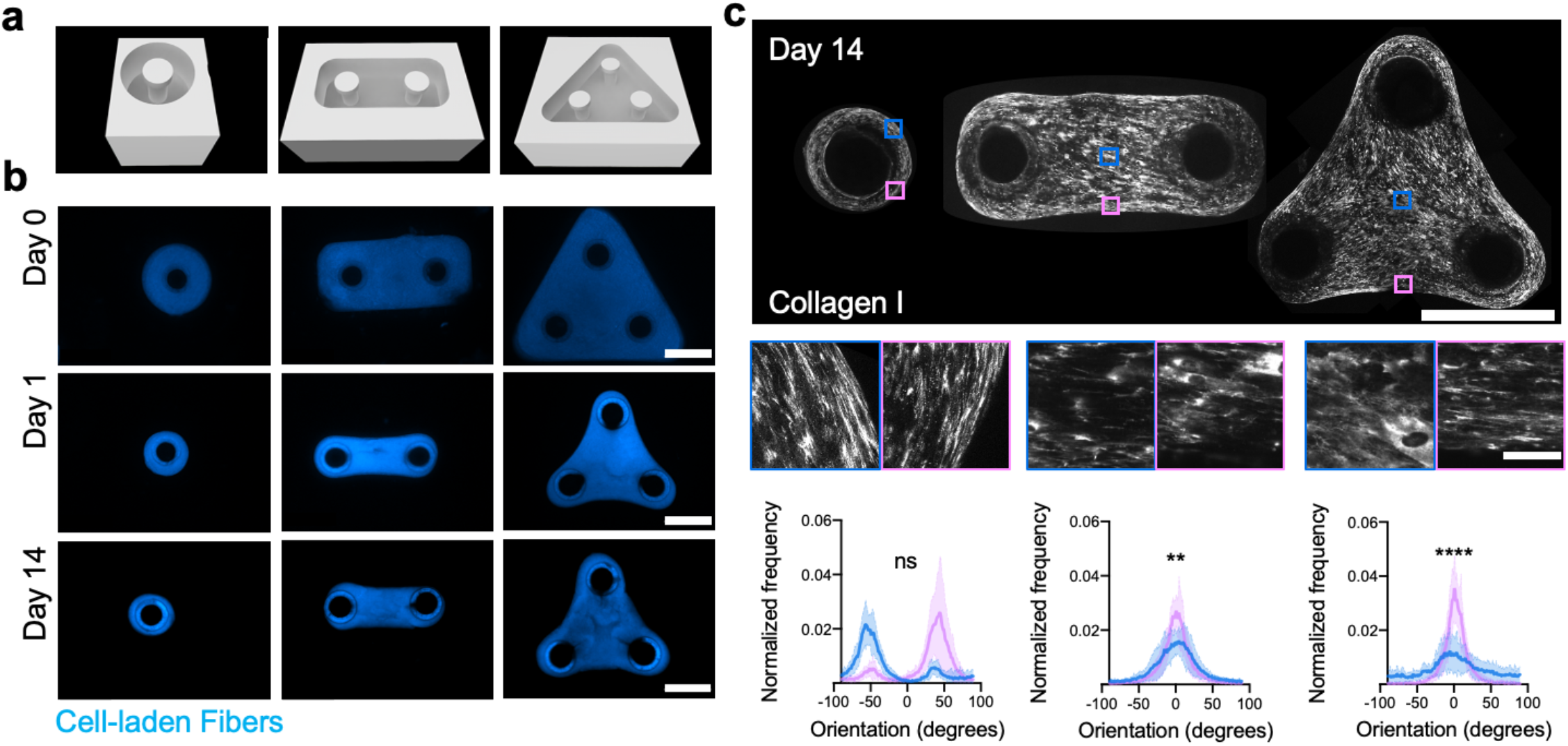
Microtissue assembly and ECM organization in fibrous hydrogel assemblies. (a) Schematic of single, two, and three post microwells used to fabricate microtissues with circumferential, uniaxial, and random alignment, respectively. (b) MSC-laden assemblies (20%, blue) formed in microwells and imaged at 0, 1, and 14 days. Images are representative of n=11, 15, 11 independent microtissues from 3 biologically independent experiments for single, two, and three post microwells, respectively. Scalebar 2 mm. (c) Representative images of collagen I labeled microtissues after 14 days of culture (top: macroscale images, scalebar 1 mm; insets: highlights of color-coded regions, scalebar 50 µm) and quantified collagen alignment profiles (bottom) from color-coded regions. (n= 11, 15, 11 independent microtissues from 3 biologically independent experiments for single, two, and three post microwells, respectively, mean ± s.d., Watson-Wheeler test for homogeneity, two post (bottom center) *p* = 0.0014, three post (bottom right) *p* = 6.4 x 10^−10^). (n.s. not significant, ** p<0.01, ****p<0.0001)

Other synthetic materials that support microtissues have been described; however, these systems utilize either covalently crosslinked gels that require cell-mediated degradation that slows down tissue formation (*48*), or hydrogels crosslinked with dynamic covalent crosslinks that rapidly erode and limit control over material properties (*49*). Alternatively, fibrous assemblies provide a material system that promotes rapid tissue formation, and materials are integrated into the final tissue structure. The ease of tuning contractility should allow for a broad range of cell types to be used across various emerging microtissue technologies (*47*). Beyond microtissue formation, fibrous assemblies could potentially be useful for other bottom-up approaches such as the self-assembly of vasculature (*2*), organoid culture and fusion (*50*), and microphysiological systems (*43, 51*).

### Programmable materials through extrusion printing and photopatterning

As the fibrous assemblies have many important features amenable to fabrication techniques, such as shear-thinning and self-healing behaviors and photocrosslinking, we explore their potential in contractile and dynamic constructs. Due to the extremely soft properties of fiber assemblies prior to photocrosslinking, we utilize an equally soft (G’< 10 Pa) support bath for the extrusion printing of low fiber density (20%) fiber solutions that would otherwise collapse if extruded onto a surface. Specifically, the support bath is composed of an agarose microgel slurry and unmodified hyaluronic acid, which allows yield stress properties and increased viscosity for handling, respectively (Supplemental Fig 14). Cell-laden fibrous assemblies (75% and 20%) are extruded into the support bath, crosslinked with light, cultured for 24 hours, released from the support, and then imaged to assess changes in filament shape and resolution (Fig 6a). High fiber density (75%) filaments do not contract, while low fiber density (20%) filaments contract down to ∼46% of their original area and increase cell density (Fig 6b), effectively increasing printing resolution.

**Figure 6.**
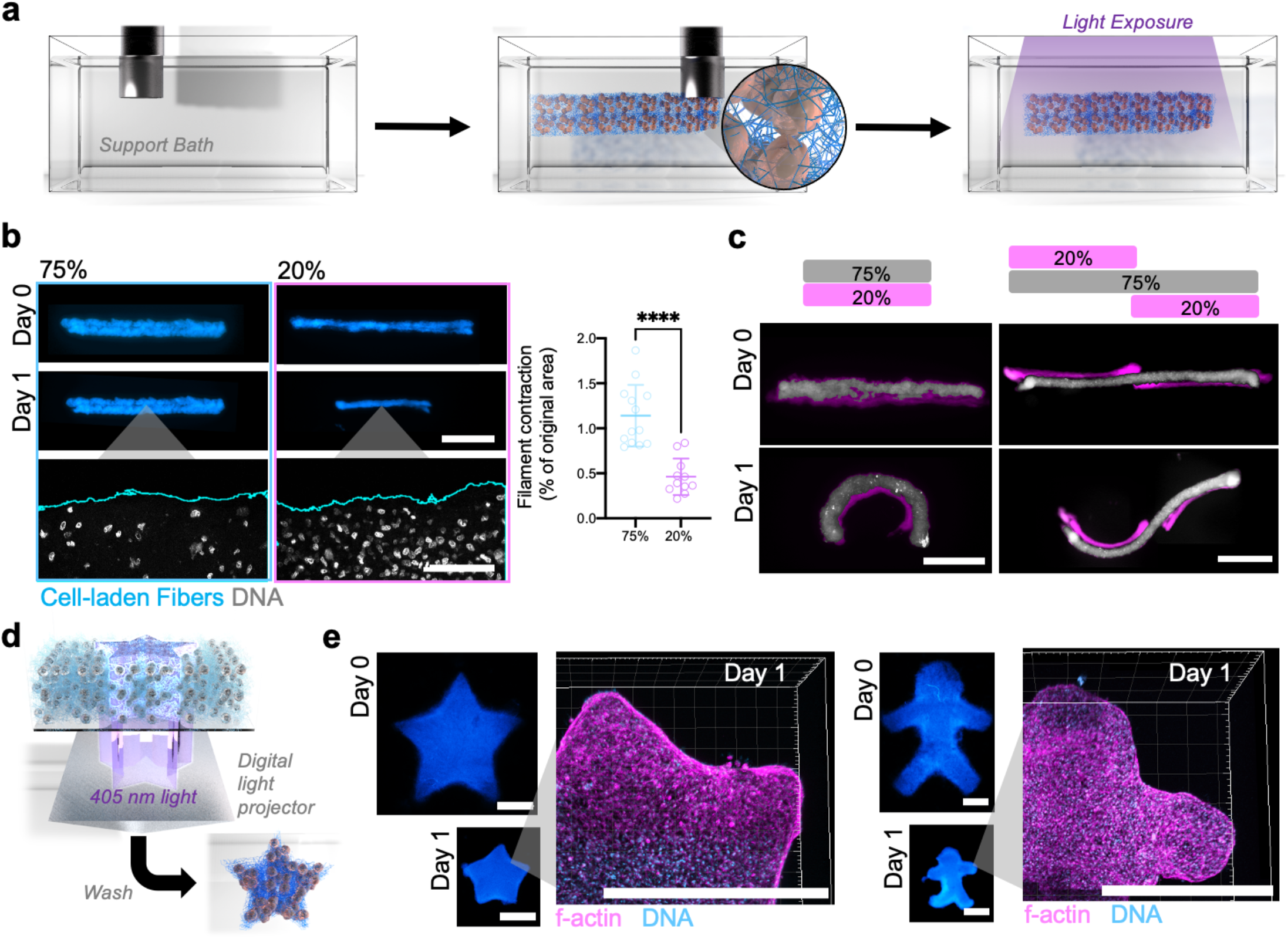
Biofabrication of contractile and programmable cell-laden structures from fibrous hydrogel assemblies. **(a)** Schematic of extrusion bioprinting of cell-laden assemblies into support hydrogel bath and photocrosslinking into stabilized structures (right). **(b)** Representative images (top: scalebar 1mm, bottom: scalebar 100 µm, filaments (blue), DNA (grey)) and quantification of bioprinted cell-laden filaments immediately (Day 0) and after culture (Day 1) (n=13, and 11 individual filaments from 3 biological independent experiments for 75% and 20%, respectively, mean ± s.d., two-tailed t-test *p* = 9.0 x 10^−6^). **(c)** Schematic (top) and representative images (bottom) of composite cell-laden filaments composed of 75% (grey) and 20% (pink) imaged immediately (Day 0) and after culture (Day 1). Images are representative of n=3 (left, single bend) and n=2 (right, multi-bend) biologically independent experiments each with 2 printed constructs. Scalebars 1 mm (single bending, left), 2 mm (multi-bend, right). **(d)** Schematic of projector-based photopatterning process (405 nm light), where structures are obtained with light-mediated patterning and washing of uncrosslinked fibers and cells. **(e)** Representative images of photopatterned star (left) and stick-figure (right) structures contracting over 1 day and staining (actin (pink), DNA (blue) after 1 day of culture. Images are representative of 3 biologically independent experiments each. Scalebar 1 mm. (****p<0.0001).

Having shown that we can print contractile and non-contractile filaments, we next explore how the fusion of 2 filaments with differential contractility might contribute to geometric changes over time. Using multi-material extrusion printing into support baths, contractile (low fiber density) and non-contractile (high fiber density) filaments are printed side by side and observed to undergo bending towards the direction of the contracting filament (Fig 6c, left). Instabilities created by differential cell contraction are hypothesized to promote tissue bending during development and in engineered tissues (*8, 52*), and our results suggest that fiber density can influence morphological changes in tissue shape. To further showcase the utility of this platform, we show that multiple bending instabilities can be introduced by printing multiple contractile filaments next to one larger non-contractile filament (Fig 6c, right). These results indicate that the system is a highly tunable method to print tissues with predictable shape changes over time.

Extrusion bioprinting resolution is generally limited to ∼100 µm, whereas emerging photolithography-based printing methods have higher resolutions with features around 10-100 µm (*53*); however, there is still much room for improvement in print resolution and with materials that mimic the ECM (*53*). Fibrous assemblies require photocrosslinking to stabilize fibers, making them ideal resins for photolithography-based printing. As a proof of concept, we use a digital micromirror device to create patterns of light to crosslink cell-laden fiber assemblies (Fig 6d), and after photocrosslinking and washing, we obtain high fidelity patterned structures (Fig 6e, Supplemental Fig 15). Star and stick-figure shaped constructs are printed and allowed to contract overnight, and a high level of contraction is observed while retaining the original printed shape and simultaneously reaching tissue level cell densities (Fig 6e). To our knowledge, these are the first photocrosslinkable contractile bioinks that have been described, and they could find utility for fabricating constructs with tissue relevant cell densities, high-resolution structures, and dynamic behaviors.

## Discussion

Advances in materials and processing techniques have enabled the development of hydrogels that capture many properties (e.g., structural, biochemical, mechanical) of natural ECMs to mimic the complexity of native tissue environments to study cell-ECM interactions, enable the development of functional model systems, and engineer tissues. Many of these technologies rely on encapsulating cells within environments that lack the microscale structure and dynamic cell responsive nature of the ECM, making it difficult for individual cells or collectives to undergo self-assembly processes that build the structures that give rise to tissue function. Fibrous assemblies allow cells to physically remodel their microenvironment to rapidly assemble organized microtissues, while maintaining control over the microscale properties of the ECM. By modulating fibrous assembly porosity, cell-induced contractility of materials can be tuned and we demonstrate the potential for fabricating high cell density aligned microtissues. We believe this to be a significant advance in the development of synthetic fibrous matrices to mimic natural ECM.

The potential application of fibrous assemblies in biofabrication techniques is explored by printing tissues that shrink to improve resolution and have programmable bending in 3D. A major goal in the biofabrication field is to create tissues that undergo morphological changes over time to mimic developmental processes or yield functional tissues (*53*–*55*). Our findings highlight how the microscale architecture and force responsive properties of the ECM influence macroscopic changes in tissue shape over time, and provide methods to program tissue bending. This was only possible through the development of new fibrous assemblies. Cell patterning (*56*), support gel fabrication (*5*), and high cell density printing (*51, 57*) are other emerging technologies to assemble tissues through cell-induced shape changes; however, most of these methods require natural ECMs that are permissive to cell interactions and function. Fibrous assemblies could be seamlessly integrated with these emerging methods, to provide a high level of control over the microenvironment and assembly process. Additionally, we believe that fibrous assemblies could enable developments in other areas, such as the fabrication of biological machines (*58*) and multi-tissue assembly. The approach is highly generalizable and could be implemented with a wide range of materials and cell types, which opens up the exciting opportunity to direct self-assembly processes such as contraction, cell differentiation, and tissue morphogenesis over time.

Although we have demonstrated that fibrous assemblies recapitulate dynamic properties of natural ECM and the importance of fiber density on 3D cell contraction and macroscale morphological changes, there remain limitations to this study. There are endless formulations that can be fabricated with respect to material composition, which will influence network degradability and crosslink stability (e.g., covalent versus physical) and the extent to which the assemblies mimic natural ECM fibers (*10*); however, we have only selected a range of material formulations for this proof-of-concept illustration of the technique and potential utility of the approach. Additionally, the impact of fiber density on contraction was the only material property investigated here, and future work should explore how other parameters such as fiber length, stiffness, and biochemical composition influence contractility and other outcomes. Lastly, mesenchymal stromal cells were the only cells utilized in this study, and future work will need to explore the bioactivity of other cell types within these fibrous assemblies, particularly with specific applications in mind. For example, the behavior of endothelial cells, epithelial cells, and multicellular systems (spheroids or organoids) that undergo self-assembly processes would be of great interest.

## Materials and Methods

### Polymer synthesis

All chemicals were purchased from MilliporeSigma unless noted otherwise. Norbornene modified hyaluronic acid (NorHA) was synthesized as previously described (*59*). Briefly, sodium HA (Lifecore) was dissolved in DI H_2_O at 2 wt % with Dowex® resin (50Wx8 ion-change resin) at a ratio of 3:1 (resin: HA by weight) and mixed for at least 30 minutes. The resin was filtered, the filtrate was titrated to pH 7.03 with diluted tetrabutylammonium hydroxide (TBA-OH), and then frozen and lyophilized. TBA modification of HA was confirmed with ^1^H NMR. HA-TBA was dissolved in anhydrous DMSO at 2 wt% with 4-(dimethylamino)pyridine (DMAP) (1.5 molar ratio to HA-TBA repeat units) and 5-norbornene-2-carboxylic acid (3:1 molar ratio to HA-TBA repeat units) under a nitrogen atmosphere. Once fully dissolved, di-tert-butyl dicarbonate (Boc_2_O, 0.4 M ratio to HA-TBA units) was injected into the vessel and the reaction was carried out for ∼20 hrs at 45 °C. The reaction was quenched with 2X cold (4°C) DI H_2_O and dialyzed against water with 0.25g NaCl/ L of DI H2O for 3 days. NorHA was then mixed with NaCl (1g NaCl/ 100ml solution) and precipitated in ice-cold acetone (1L acetone/100ml solution). The precipitate was re-dissolved in DI H_2_O, dialyzed for 5 days, frozen, and lyophilized. Norbornene functionalization was confirmed with ^1^H NMR (Supplemental Fig1).

Norbornene modified gelatin (NorGel) was synthesized as previously described (*60*). Briefly, 10 wt% gelatin was diluted in DPBS and heated to 60 °C until fully dissolved, pH was adjusted to 7.5, 20 wt % carbic anhydride was added dropwise to the solution while stirring, and the reaction was carried out for 3 hours while maintaining Ph ∼8 with sodium hydroxide. The reaction was quenched using 5X volume of preheated DPBS, stirred for 15 minutes at 60 °C, dialyzed at 40°C for 7 days, frozen and lyophilized. Methacrylated HA (MeHA) was synthesized as previously described (*61*). Briefly, 2 grams of HA were added to 200 ml of DI H_2_O to achieve a 1wt% solution of HA. Next, while maintaining a pH of 8.5-9, 1.5 ml of methacrylic anhydride was added dropwise and mixed on ice for 8 hours. The solution was mixed for an additional 12 hours at room temperature without maintaining pH, dialyzed for 7 days, frozen, and lyophilized. Norbornene modified PEG (NorPEG, 8-arm PEG-Norbornene, 20 kDa) and thiol modified PEG (4-arm PEG thiol, 5 kDa) were purchased from JenKem.

### Electrospun fiber mat preparation

To electrospin NorHA, 3.5 wt% NorHA, 2.5 wt% PEO (900 kDa), 0.05 (v/v) % Irgacure 2959, and 4 mg/ml fluorescent dextran (FITC-dextran, 2 MDa) were mixed with various stoichiometric ratios of dithiothreitol (DTT) to norbornene groups (i.e., 0.1, 0.25. 0.45 thiols:norbornenes) in DI H_2_O, mixed at 250 rpm for 24 hours protected from light, and loaded into syringes for electrospinning. To electrospin NorGel, 5 wt% NorGel, 3.5 wt% PEO (900 kDa), 0.05 (v/v) % Irgacure 2959, and 4 mg/ml fluorescent dextran (FITC-dextran, 2 MDa) was mixed with a 3.75 mM 4-arm PEG-thiol solution in DI H_2_O. To electrospin NorPEG, 5 wt% 8-arm PEG-norbornene, 10 wt% 4-arm PEG-thiol, 2.5 wt% PEO (900 kDa), 0.05 (v/v) % Irgacure 2959, and 4 mg/ml fluorescent dextran (FITC-dextran, 2 MDa) was mixed in DI H_2_O. To electrospin MeHA, 3 wt% (MeHA), 2 wt% PEO (900 kDa), 0.05 (v/v) % Irgacure 2959, and 4 mg/ml fluorescent dextran (FITC-dextran, 2 MDa) was mixed in DI H_2_O.

For single polymer electrospinning, the electrospinning jet was positioned 19 cm away from a rotating collecting mandrel, with a flow rate of 0.7 ml/hr and voltage +28-30 kV were used. For multi-fiber electrospinning of stable (NorHA) and sacrificial (PEO) fiber populations, electrospinning jets were positioned on opposite sides of the electrospinning mandrel (Supplemental Fig 5), while maintaining the same needle to collector distance (19 cm), and flow rates were modulated (1.05 ml/hr (PEO), 0.7 ml/hr (NorHA)) to give a 60:40 ratio of sacrificial to stable fibers within the fiber mat. Electrospinning was carried out in a custom humidity-controlled chamber (15-21% humidity) with a rotating mandrel (−5kV, ∼350 RPM), and needle gauge of 18. All fiber mats derived from norbornene modified polymers were photocrosslinked for 1 hour on each side of the fiber mat surface at 10 mW/cm^2^ with UV light (320-390 nm, Omnicure S1500 UV, Spot Cure Systems), while MeHA was photocrosslinked for 15 minutes on each side of the fiber mat surface at 15 mW/cm^2^ with UV light.

### Cell isolation and culture

Human MSCs were isolated from bone marrow aspirate (Lonza) as previously described (*62*). Briefly, bone marrow was diluted with DPBS (1:4) and cells were separated with a Ficoll gradient (800 x G for 20 minutes). Mononuclear cells were isolated from the liquid interface and seeded on tissue culture plastic, and cultured in MSC expansion medium (α-MEM, 10% FBS (Gibco), 1% penicillin/streptomycin, 5 ng/ml basic fibroblast growth factor) at 37 °C until reaching 80% confluency. Cells were then trypsinized and stored in liquid nitrogen. All MSCs were passaged one to three times in MSC culture medium (α-MEM, 10% FBS, 1% penicillin/streptomycin), trypsinized using 0.05 %Trypsin/EDTA (Gibco), and resuspended in DPBS before encapsulation. All assemblies were cultured in MSC culture medium and the medium was replaced every day.

### Fiber fragmentation and purification

8 mg of electrospun fiber mats were segmented (multiple ∼2×2 mm squares), hydrated with 6 ml of DPBS in a sterile scintillation vial for 10 minutes, and then ∼3ml of the solution was repeatedly (40x) and rapidly aspirated and extruded through an 18G needle (BD) in the same scintillation vial using a 3 ml syringe. A 21G needle was then used to further fragment fibers with 40 repeated steps of aspiration and extrusion. Further fragmentation was achieved by using this same method with a 23G needle, 25G needle, 27G needle, and last a 30G needle. Fiber solutions were purified by passing the solution sequentially through a 40 µm cell strainer (Pluriselect) and then a 5 µm cell strainer. Fibers were then pelleted with centrifugation at 10,000 x G, resuspended, and washed with 1 ml of PBS, which was repeated twice to remove excess DTT crosslinker and I2959 photoinitiator. The fiber pellet volume was then measured by centrifuging, removing the supernatant completely, and resuspending in a known volume. The volume displaced by the pellet was taken as the volume of fibers at a 100% volume fraction. Fiber solutions were sterilized with germicidal UV and stored at 4**°**C.

### Fiber assembly and cell encapsulation

Various concentrations of fibrous hydrogel assemblies (20%, 50%, and 75%, v/v%) were formed by mixing the concentrated fiber solution with various amounts of PEG-thiol crosslinker (4-arm PEG thiol, 5 kDa) based on fiber volume concentration (0.1 mM for 20%, 0.25 mM for 50%, 0.375 mM for 75%), 0.05 wt% photoinitiator (lithium phenyl-2,4,6-trimethylbenzoylphosphinate (LAP), Colorado Photopolymer Solutions (Boulder, CO)), in DPBS. For cell encapsulation, these components were first mixed and cells were added last and kept on ice until use. Fibrous hydrogel assemblies were photocrosslinked using visible light (400-500 nm, Omnicure S1000) at an intensity of 5 mW/cm^2^ for 5 minutes. Cell-laden constructs were carefully washed with cell culture media immediately after crosslinking, and media was changed 30 minutes later to remove any residual radicals and photoinitiator. Microfabricated polydimethylsiloxane (PDMS) (SYLGARD 184, DOW) wells (2 mm diameter) and 0.5 ml cut syringes were used as molds for low cell density and high cell density contraction assays, respectively.

### Material and cell imaging analysis

To measure fiber lengths, a custom FIJI macro was written to binarize fluorescent images of dilute fiber solutions, and ridge detector was used to measure fiber lengths. Fiber diameter and porosity were measured using previously described methods (*61*). Void fractions of fibrous hydrogel assemblies were measured by thresholding fluorescent images of assemblies, inverting the images, and measuring the % area of the signal. This gives the void fraction for a xy plane by measuring the area of the pores, and this measurement was used to measure the average void fraction for 10 slices through ∼10 µm in the z-direction.

Cell shape descriptors were obtained from max z-projections of phalloidin and Hoechst 33342 stained cells. Images were binarized and aspect ratios and cell circularity were measured using FIJI. To measure fiber compaction around cells, a region of interest (ROI) was created around cells on a single xy plane near the z position of the cell nucleus, and fiber fluorescence intensity was measured in 1 µm band increments away from the cell border (Supplemental Fig 8). Fiber fluorescence intensity around cells was compared to a remote location without cells and normalized by the average fluorescence intensity (+ s.d.) for remote locations. Cell viability was measured by casting assemblies in wells at initial timepoints or by sectioning gels with a scalpel and staining with Hoechst 33342 (5 µg/ml, stains all cell nuclei, Fisher Scientific) and ethidium homodimer-1 (4 µM, stains dead cell nuclei, Invitrogen) for later timepoints.

Macroscale contraction of assemblies was measured by manually segmenting and measuring the construct area from phase-contrast images over time. Volumetric changes were obtained by measuring the height and circumference of contracted assemblies after 3 days of culture using phase-contrast microscopy. Contraction of microtissues, extrusion printed constructs, and photopatterned constructs were imaged with fluorescence and brightfield microscopy.

### Rheological and Mechanical characterization

The viscoelastic, photocrosslinking, and stiffening properties of fiber solutions, fibrous hydrogel assemblies, and support hydrogels were characterized using shear rheology (cone and plate geometry with (0.995°) cone angle and a 27 µm gap, or parallel plate geometry, TA Instruments, AR2000) at room temperature. To assess the shear-thinning properties, the storage modulus (G’) and viscosity were measured as a function of strain (0 to 5) and shear rate (0 to 50/s), respectively. Photocrosslinking of fibrous hydrogel assemblies was measured in real-time at low strain (0.01) at 1 Hz. To assess strain-induced stiffening properties, G’ was measured at a function of increasing strain (0.005-1) at 1 Hz. Self-healing of fiber solutions and support gels were assessed by applying low (0.01) and high (5) strains repeatedly at 1 Hz. The compressive moduli of contracted assemblies were measured using dynamic mechanical analysis (TA Instruments, Q800).

### Microwell fabrication, microtissue formation, and photopatterning

Microwell hydrogels with various post geometries were fabricated using a DLP printer (Volumetric, LumenX) with PEG-diacrylate resin (PEGDA PHOTOINK TM), and MSC-laden fiber solutions were seeded into microwells with a 20 µl pipette, and then crosslinked with light. Microtissues were then hydrated with MSC culture medium and the medium was changed 30 minutes later and every day after for 14 days.

To photopattern cell-laden structures, custom PDMS wells were fabricated and cell-laden fiber solutions were added to the well. The DLP printer was used to project various photopatterns (i.e., star and stick figure) that photocrosslinked the fibrous hydrogel assemblies along the light path and created stable 3D structures. A light intensity of 20 mW/cm^2^ (405nm LED) for 30 seconds was used to create photopatterned constructs. After photopatterning, constructs were gently hydrated with small droplets of MSC culture medium near the edge of the print and then flooded with excess media. Constructs were imaged immediately after washing and after 1 day of contraction.

### Support gel preparation and extrusion bioprinting

Agarose support gels were made as previously described (*63*), with some modification. Briefly, a 0.5 wt% agarose (SeaKem, Lonza) solution was mixed and autoclaved to melt and sterilize, and then cooled while stirring at 700 rpm to form microgels. This microgel solution was then diluted in an 8 wt% solution of HA, to give a final agarose concentration of 0.25 wt% and HA concentration of 4 wt%. Agarose was diluted to give a support gel that was soft enough to yield with extremely soft fiber solutions. HA was added to diluted agarose microgel solution to increase the viscosity and stabilize the printed structures while moving. Fiber solutions were extruded using 27G needles and a sub microliter injection needle (World Precision Instruments, Nano-Fil 100), with two different extrusion printers (Allevi 2, and Velleman K8200), and photocrosslinked (400-500 nm, 5 mW/cm^2^ for 5 minutes) after printing. Filaments were imaged directly after printing, as well as after releasing prints from the support bath at ∼20 hours of culture by diluting with an equal volume of MSC media.

### Immunofluorescent labeling and staining

After terminal timepoints, cultures were fixed with pre-warmed 4% PFA, for 30 minutes, and then washed with DPBS 3x. Large constructs used in macroscale contraction assays were fixed overnight at 4°C, cryoprotected in 30% sucrose solution, embedded in OCT compound, flash-frozen by dipping into liquid nitrogen cooled 2-methyl butane, and cryosectioned (20 µm sections). Actin and DNA were visualized by staining with Alexa fluor-647 phalloidin (Cell Signaling Technology) and Hoechst 33342 (Fisher Scientific). For immunolabeling, sections or microtissues were hydrated, blocked with 1% BSA in DPBS for 1 hour at RT, and then incubated with either anti-fibronectin (1:50, Sigma F6140) or anti-collagen type I (1:200, Abcam ab138492) in 1% BSA buffer overnight at 4°C. The next day, sections and microtissues were washed 3x with DPBS, and then incubated with the secondary antibodies Alexa Fluor 647 anti-mouse (1:200, A-21235, Fisher Scientific) and Alexa Fluor 594 anti-goat (ab150080) for two hours at room temperature, and then counterstained with Hoechst 33342. Sections and microtissues were imaged using a Leica SP5 confocal microscope.

### Fiber network model

Three-dimensional discrete Voronoi fiber networks were employed to model the networks (*37*). To generate a Voronoi network, first, random seed points were chosen in a three-dimensional domain. The Voronoi diagram corresponding to these seed points was then generated using MATLAB (MathWorks, Natick, MA). Beam elements, modeling fibers were considered along the edges of the resulting Voronoi diagram. To perform mechanical tests, we extracted a cubic sample from the constructed network. The fibers were curved to mimic the experimental images and also facilitate the computations. Individual fibers were modeled as elastic rods with circular sections, capable of bending, stretching, and shear. Hybrid beam formulation and reduced integration were used to facilitate computations. The fiber elastic modulus was 1 MPa.

In this model, the dimensionless ratio of fiber diameter to Voronoi cell diameter controls the network’s deformation. The variations of network density were modeled using fiber to cell diameter ratios of about 1%, 4%, and 6%, which were obtained by iteration through various fiber to cell diameter ratios until these network densities matched the experimental data with 20%, 50% and 75% fibrous hydrogel assemblies, respectively. The mean connectivity (nodal coordination) of the network model is between 3 and 4, to match the experimentally observed network connectively. Displacement-controlled tests were performed by prescribing the displacement of boundary nodes. Shear tests were performed by fixing the displacement of the nodes at the bottom surface of the network while horizontally moving the nodes at the top surface. Additionally, the networks were stretched in uniaxial tests while free to contract in the directions transverse to loading. Implicit finite element calculations were performed using the ABAQUS software package (Simulia).

### Statistical analysis

Microsoft Excel was used to store raw data, while GraphPad Prism 9 was used for all statistical analysis. Comparisons between two experimental groups were performed using two-tailed t-tests and comparisons among more groups were performed using a one-way ANOVA with Tukey post hoc testing or multiple groups over time using either a two-way ANOVA with Tukey post hoc test or a mixed-effects analysis with Sídák post hoc testing. Circular statistics were performed using the package “circular” in R software, where Watson-Wheeler test for homogeneity was carried out on 2 different groups (*64*). All n numbers refer to biologically independent samples with no repeated measures and p-values are provided in figure legends. The sample distributions were assumed to be normal with equal variance.

## Acknowledgements

We acknowledge Dr. Claudia Loebel, Dr. Kwang Hoon Song, Dr. Leo Wang, and Victoria Muir for help and advice with MSC isolation and culture, polymer synthesis, material fabrication, and NMR spectroscopy, respectively.

## Author contributions

M.D. and J.B. conceived the ideas, designed the experiments. M.D., M.P., E.B, K.X., G.M., P.M., A.D., conducted experiments and computational studies and interpreted the data. M.D. and J.B. wrote the manuscript. M.D., M.P., E.B, K.X., G.M., P.M., A.D., P.J., V.S., and J.B. interpreted the data and edited the manuscript. All authors have given approval to the final version of the manuscript.

## Data and materials availability

All data needed to evaluate the conclusions in the paper are present in the paper and/or the Supplementary Materials. Additional data related to this paper may be requested from the authors.

