## Supplementary Materials for "Programmable and Contractile Materials Through Cell Encapsulation in Fibrous Hydrogel Assemblies"

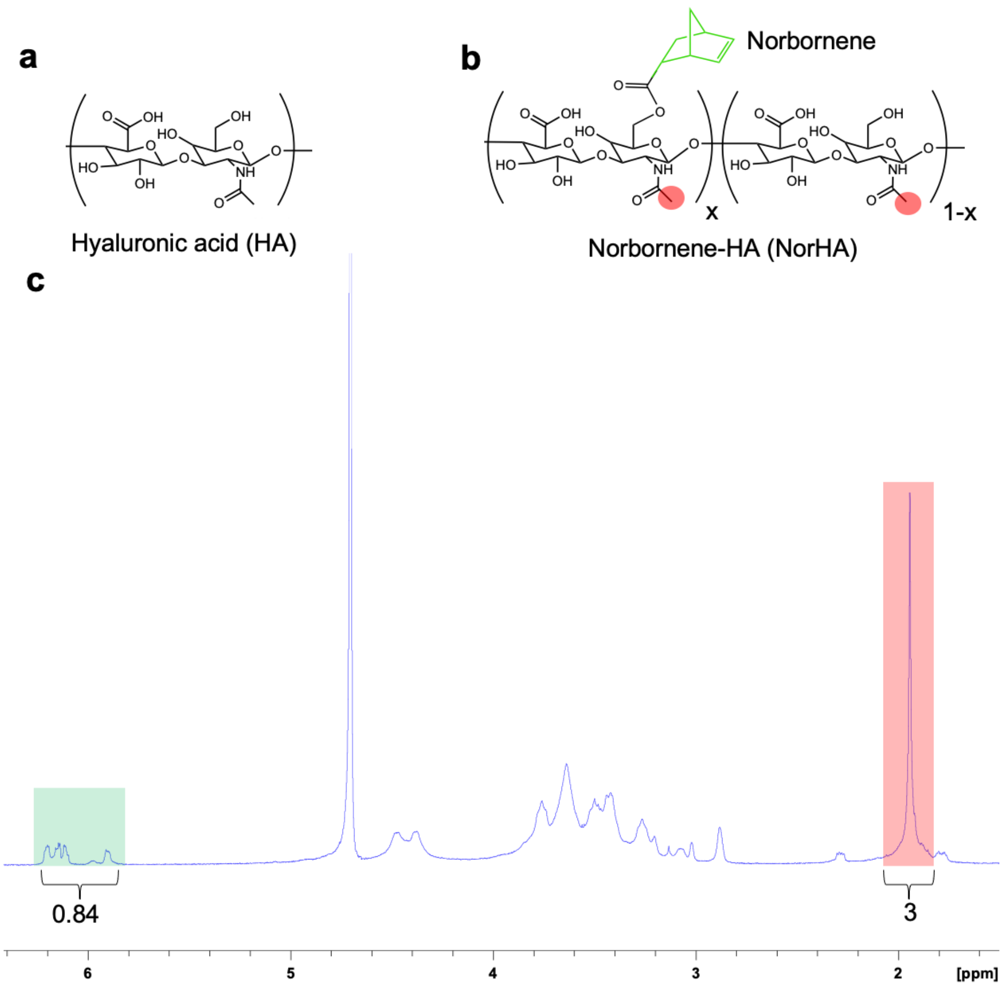


**Figure S1. ¹H NMR characterization of norbornene modified hyaluronic acid.** (a) Structure of unmodified hyaluronic acid (HA) and (b) norbornene modified HA (NorHA). (c) Norbornene modification (~40%) of HA was determined by integration of the vinyl peaks (2H, highlighted in green) relative to methyl group of HA (3H, highlighted in red).

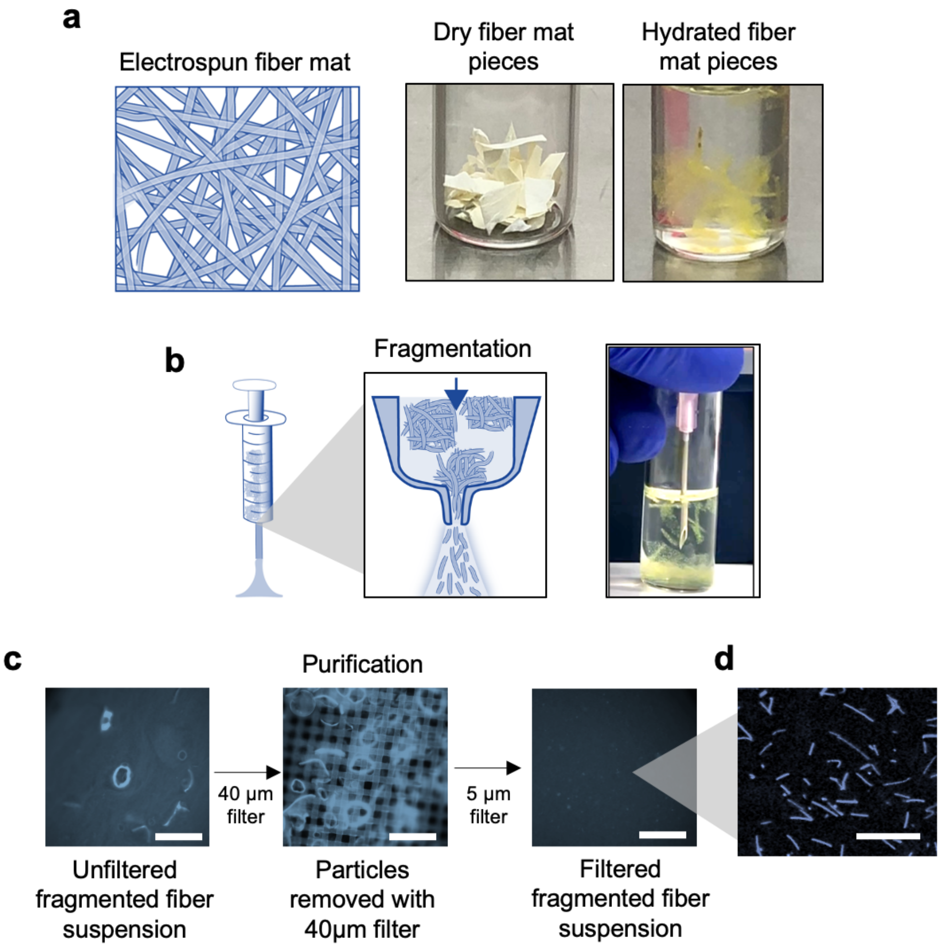


**Figure S2. Fragmentation method and isolation of fiber suspensions.** (a) Polymer solutions (e.g., NorHA) are electrospun into scaffolds using standard techniques, crosslinked, and cut into pieces that are hydrated for processing. (b) Hydrated scaffold pieces are fragmented with repeated passing (40x) through a needle (e.g., 18G) to obtain a fragmented fiber suspension. (c) Fragmented fiber suspensions retain some irregularly sized scaffold particles (left) that are filtered from the suspension by passing through a 40 μm cell strainer (center) and then a 5 μm filter (right) to obtain (d) a pure fiber population (images representative of n=4 separate electrospun fiber mats, scalebar 1 mm for (c) and 50 μm for (d)).

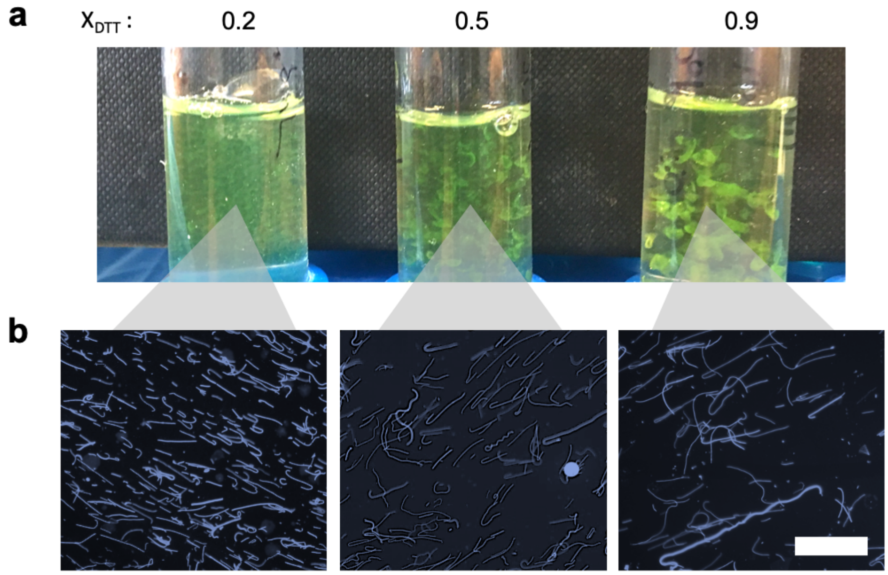


**Figure S3. Influence of the extent of fiber crosslinking on fragmentation into fiber suspensions.** (a) Digital images of hydrated electrospun fiber mat pieces composed of NorHA fibers with varied amounts of norbornene consumption ($X_{DTT}$: 0.2, 0.5, 0.9) during processing of fiber mats (n= 2 independent fragmentation experiments). (b) Representative fluorescent images of fiber suspensions after fragmentation (40x passes through 18G needle) and purification. Overall, there is increased fiber yield and uniformity and shorter fibers with lower norbornene consumption (0.2), when compared to progressively longer fibers with smaller yields with increased norbornene consumption (0.5, 0.9) (n=2 independent fragmentation experiments, scalebar 100 μm).

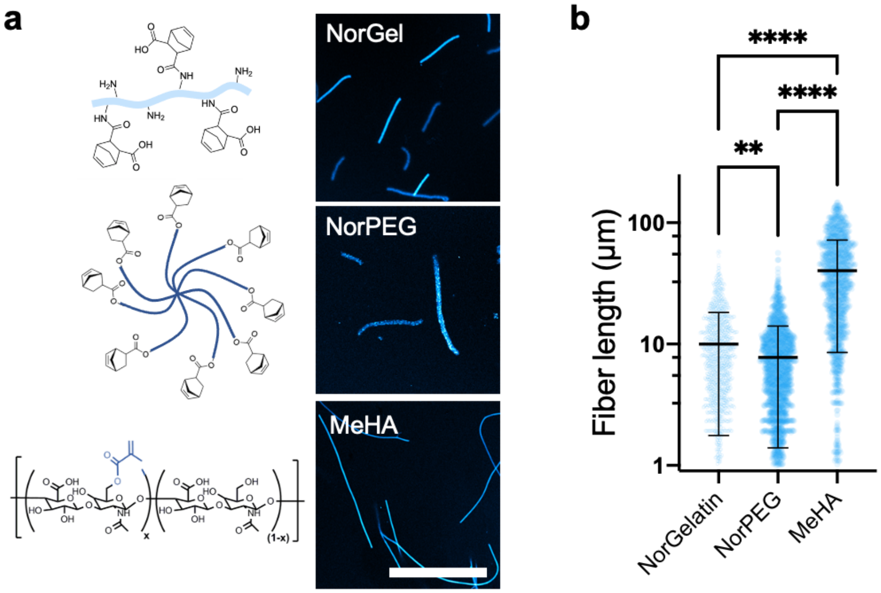


**Figure S4. Fabrication of fiber suspensions from various electrospun hydrogels.** (a) Schematics and chemical structures (left) and representative images of fiber suspensions (right) generated from the fragmentation (40x passes through 18G needle) of electrospun fiber mats composed of norbornene modified gelatin (top, NorGel), norbornene modified 8-arm poly(ethylene glycol) (middle, NorPEG) and methacrylated hyaluronic acid (bottom, MeHA) (images representative of n=2 electrospun fiber mats, scalebar 50 µm). (b) Quantified fiber lengths from fiber suspensions generated through the fragmentation process (mean ± s.d., NorGel: n=1213, NorPEG: n=1576, MeHA: n=1208 individual fibers measured from 2 independent fragmentation experiments, mean ± s.d., one-way ANOVA, (NorGel vs. NorPEG, p=0.004, NorGel vs. MeHA, p= 3.0 x $10^{-8}$, NorPEG vs. MeHA, p= 3.0 x $10^{-8}$)). (**$p<0.01$, ****$p<0.0001$).

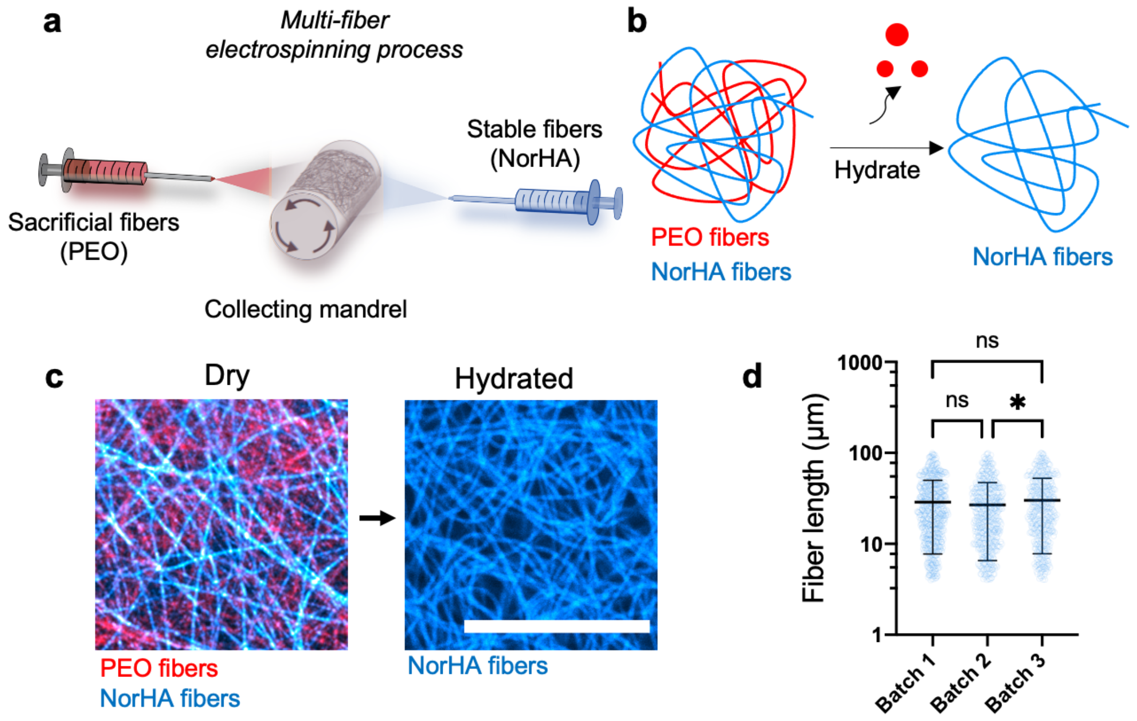


e

| | Fiber length | Fragmentation time (40x steps) | Yield (ul of fibers/mg starting material) |
|---|---|---|---|
| +Sacrificial fiber | 24 µm (+/- 11) | 5 minutes | 8 µl/mg |
| -Sacrificial fiber | 22 µm (+/- 13) | 20 minutes | 2 µl/mg |

**Figure S5. Sacrificial multi-fiber electrospinning to improve fiber yield and processing time.** (a) Schematic of multi-fiber electrospinning where poly(ethylene oxide) (PEO, red) is the sacrificial fiber population and NorHA (blue) is the stable fiber population collected on a rotating mandrel. The fiber ratio (60:40, sacrificial:stable) is tuned with flow rate of electrospinning solutions (1.05 ml/hr: 0.7 ml/hr). (b) Schematic of the washing of PEO fibers with hydration to retain only NorHA fibers. (c) Representative images of multi-fiber electrospun mats before and after hydration and washing of PEO fibers (images representative of n=3 separate electrospun fiber mats, scalebar 25 µm). (d) Measure of fibers across 3 independent batches of fiber suspensions to illustrate intra-experimental variability of the fragmentation process (40x passes through 18G needle) (Batch 1: n=674, Batch 2: n=563, Batch 3: n=490 individual fibers, mean ± s.d., one-way ANOVA, (Batch 2 vs. Batch 3 p=0.03)). (e) Table of fiber length, fragmentation time, and yield of fragmentation process comparing samples with or without the PEO sacrificial fibers when starting with 8 mg of a fiber mat (n=3 independent fragmentation experiments). (ns: not significant, *p<0.05).

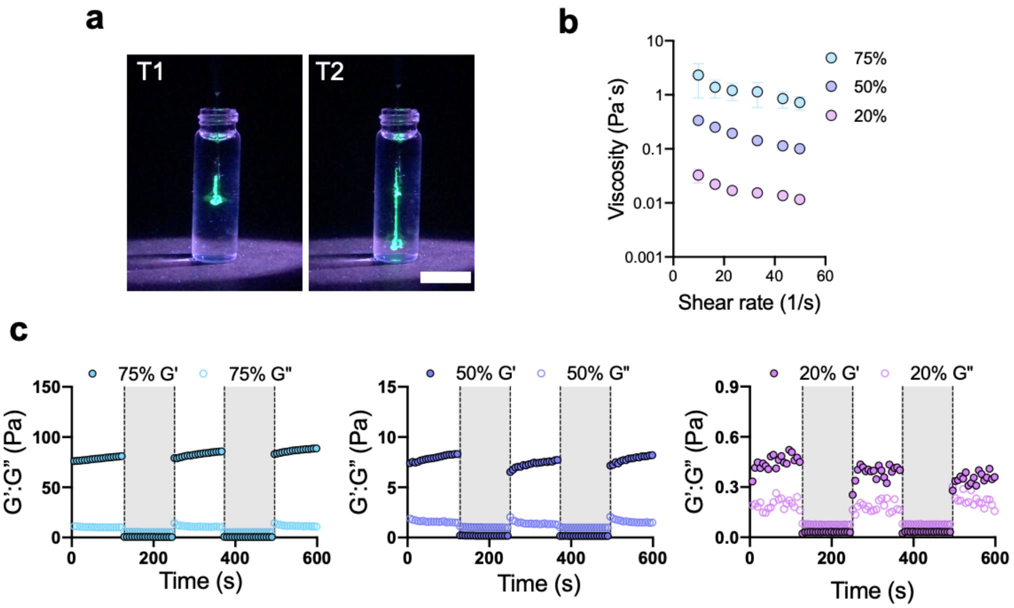


**Figure S6. Injection and rheological properties of fibrous hydrogel assemblies prior to photocrosslinking.** (a) Representative digital images of injection of FITC labeled 50% assemblies into saline at time 1 (T1, left) and time 2 (T2, right) with UV illumination for visualization (images representative of n=3 assembly batches injected, scalebar 20 mm). Rheological characterization of assemblies demonstrating (b) shear-thinning behavior as decreased viscosity with continuously increasing shear rate (0–50 $s^{-1}$) (mean ± s.d, n=3 separate samples) and (c) self-healing behavior as the recovery of storage modulus during repeated periods of low (1% strain, 1 Hz) and high (shaded, 500% strain, 1 Hz) strain cycles (n=3 separate samples).

a b

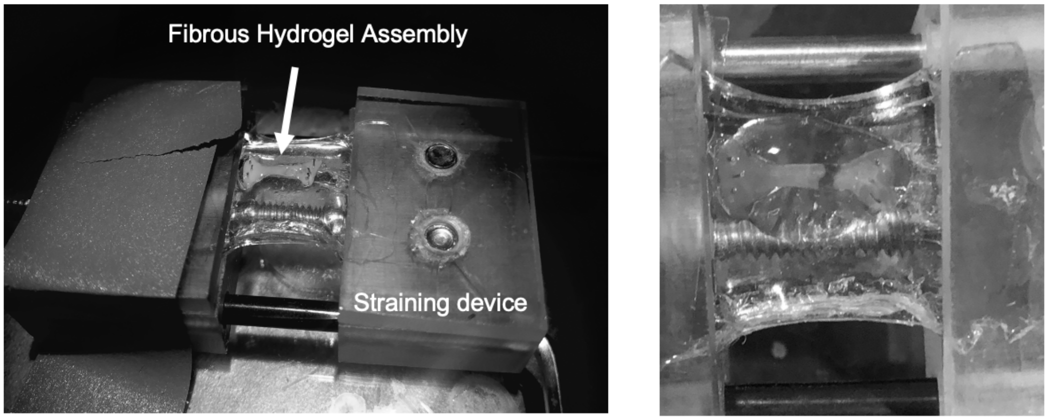


**Figure S7. Straining device and sample failure.** (a) Digital image of custom stretching device used for straining fibrous hydrogel assemblies in tension that were processed into dog-bone shapes via molds, photocrosslinked, and hydrated. Devices were placed on a confocal microscope and imaged with and without strain (10%). (b) Samples were then strained to failure and observed to fail mid-substance. Images are representative of n=3 individual samples.

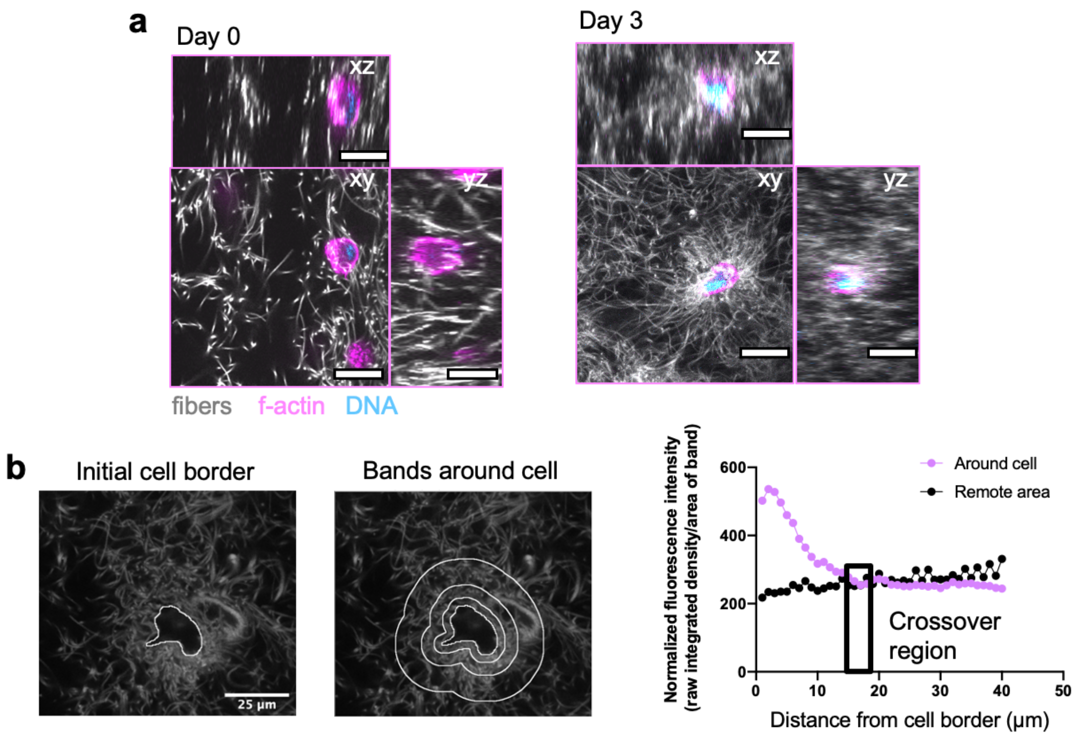


**Figure S8. 3D cell encapsulation and fiber compaction quantification methods.** (a) Orthogonal (xz, yz) and xy slices of MSCs encapsulated in fibrous hydrogel assemblies at day 0 and day 3 (n=3 biologically independent experiments, scalebar 25 µm). (b) Method for measuring fiber compaction length, including (i) 2D slices near the z position of the cell nucleus used to outline the cell and convert to regions of interest (ROIs), (ii) progressively expanded outlines of 1µm increments introduced from the cell surface and the fluorescence intensity of the channel corresponding to FITC labeled fibers measured and normalized to area (image shows example of bands at 3, 8, and 18 µm), (iii) values compared to a remote location without cells, and (iv) fiber compaction length calculated as the point where the fiber fluorescence intensity crosses the remote location fluorescence intensity plus its standard deviation.

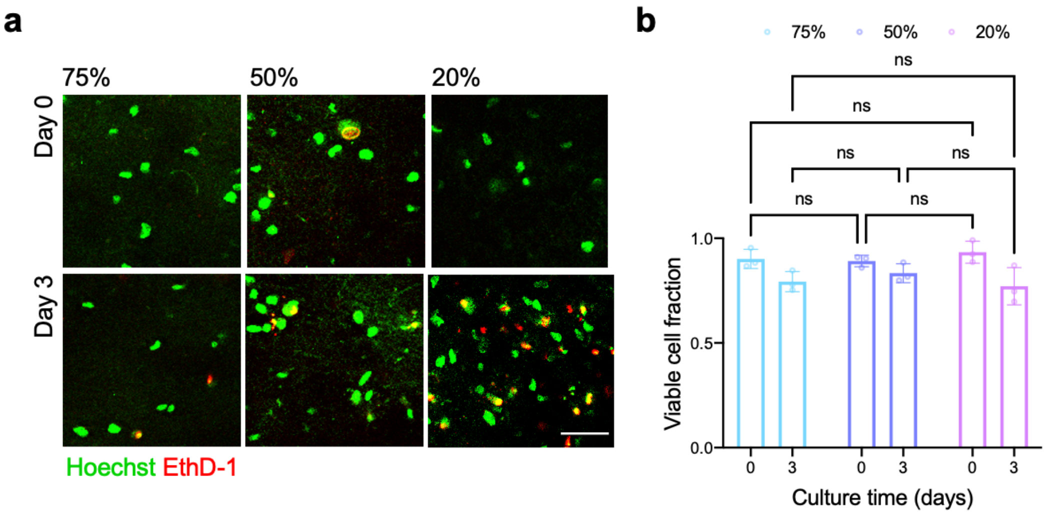


**Figure S9. Viability of cells encapsulated within fibrous hydrogel assemblies.** (a) Max z-projections (10 µm) of live/dead MSC staining 0 and 3 days after encapsulation within assemblies of varied fiber densities (75%, 50%, 20%). Hoechst 33342 (green) stains all cell nuclei while Ethidium Homodimer-1 (red) stains dead cell nuclei (n=3 biologically independent experiments, scalebar 50 µm). (b) Quantification of the fraction of viable cells after 0 and 3 days (n=3 biologically independent experiments each with 3 separate samples, mean ± s.d., two-way ANOVA). (ns: not significant).

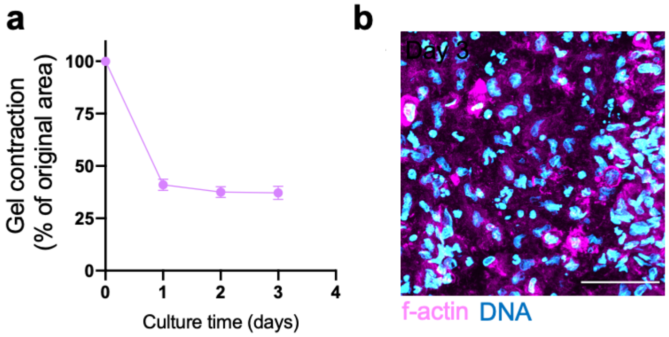


**Figure S10. Behavior of fibrous hydrogel assemblies with high densities of encapsulated MSCs.** (a) Bulk contraction of assemblies with high MSC density ($5x10^7$ cells per ml) encapsulated and measured over 3 days with phase contrast microscopy (n=3 biologically independent experiments each with ≥3 separate samples, mean ± s.d.). (b) MSC density measured after 3 days of contraction using confocal microscopy. Scalebar 50 µm.

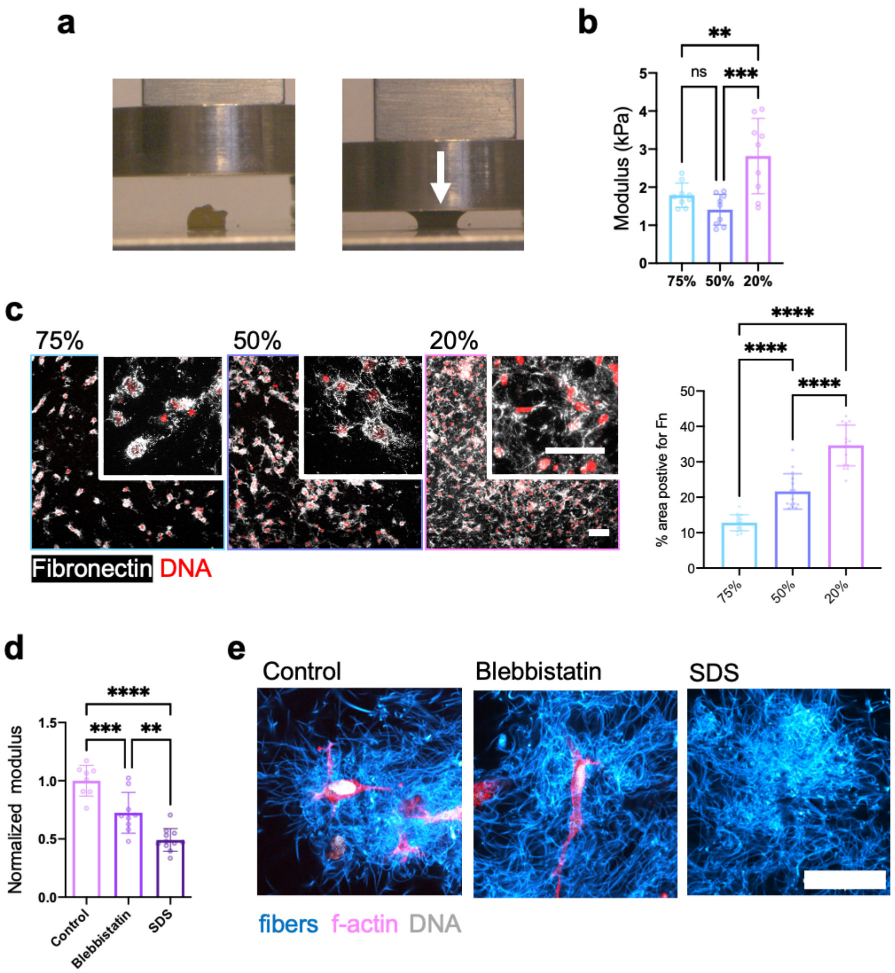


**Figure S11. Contracted fibrous hydrogel assembly mechanical properties.** (a) Images during compression testing and (b) quantified compressive modulus of assemblies across varied fiber densities (75%, 50%, 20%) with encapsulated MSCs (1x$10^6$ cells/ml) and culture for 3 days. Images and compressive moduli are representative of n= 3 biologically separate experiments, each with 3 separate samples per condition (mean ± s.d., one-way ANOVA, (20% vs. 50% $p$ = 0.0003, 20% vs. 75% $p$ = 0.007)). (c) Representative images (left) and quantification (right) of cellular fibronectin within contracted assemblies. Images and quantification are representative of n=3 biologically independent experiments, each with 3 separate samples (mean ± s.d, one-way ANOVA, (20% vs. 50% p= 3.0 x $10^{-9}$, 20% vs. 75% p= 1.0 x $10^{-12}$ , 50% vs. 75% p= 0.00001). Scalebar 50 µm. (d) Normalized modulus of assemblies from 20% fiber density after 3 days, including treatment to inhibit myosin II with acute blebbistatin or cell lysis with sodium dodecyl sulfate (SDS) (control: n=8 samples across 3 experiments, each with ≥2 samples; blebbistatin: n=9 samples across 3 experiments, each with 3 samples; SDS: n=11 samples across 3 experiments, each with ≥2 samples (mean ± s.d., one-way ANOVA, control vs. blebbistatin $p$ = 0.0009, control vs. SDS $p$ = 6.0 x $10^{-8}$, blebbistatin vs. SDS p=0.002)). (e) Representative images of control assemblies with MSCs encapsulated and cultured for 3 days, including with treatment with blebbistatin and SDS (images representative of n=2 biologically separate experiments, each with 3 separate samples, scalebar 50 µm). (ns: not significant, **p<0.001 ,***p<0.001, ****p<0.0001).

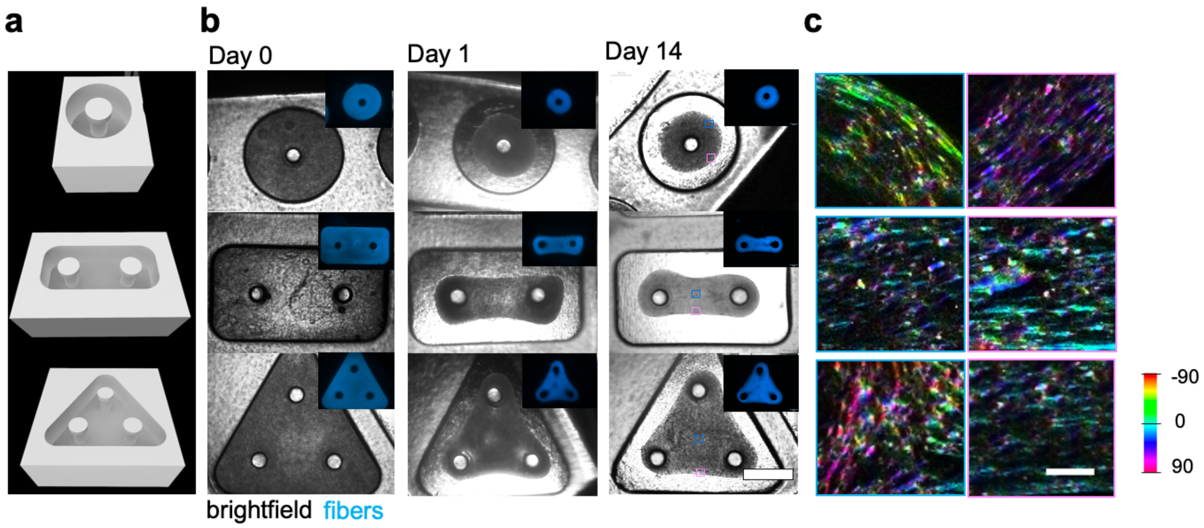


**Figure S12. Microtissue formation and ECM alignment.** (a) Digital render of stereolithography (.stl) file used for digital light processing (DLP) based printing of microtissue wells. (b) Phase contrast images of microtissues over time (black and white is brightfield, blue insets are fluorescently labeled fibers) (images representative of n= 3 biologically separate experiments, each with 3 separate samples, scalebar 1 mm). (c) Color representation of cell deposited collagen I fiber orientation within microtissues. Blue and pink squares on day 14 images in (b) correspond to approximate locations where ECM alignment was measured in (c) (images representative of n=3 biologically separate experiments, scalebar 100 µm).

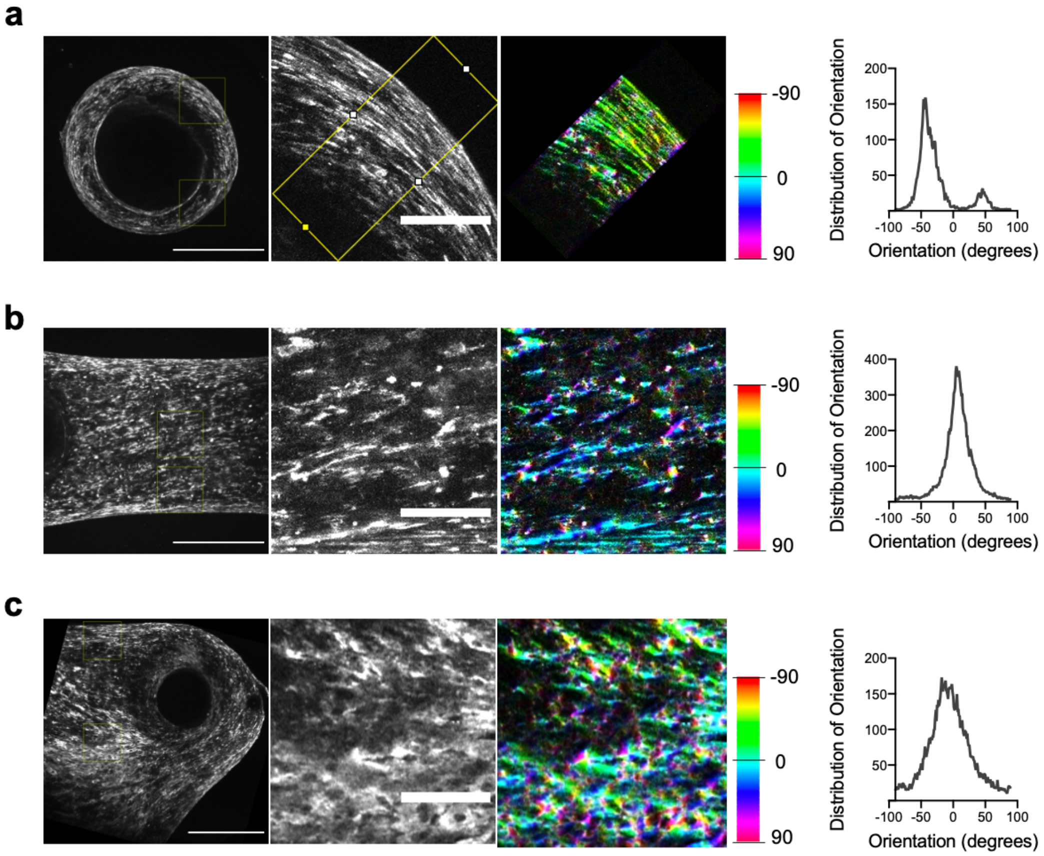


**Figure S13. Methods to quantify collagen orientation in microtissues.** (a) For single post microtissues, regions of interest (ROIs) are designated on the top and bottom of one side of a microtissue (left), each ROI is duplicated (center), and a rounded rectangle is used to make an ROI at a 45 degree angle and this ROI is used to measure fiber orientation with orientation J (right image, color represents fiber orientation, right graph is orientation distribution). (b) For two post microtissues, ROIs are made in the center and edge of the tissue as shown (left), each ROI is duplicated (center), and this ROI is used to measure fiber orientation with orientation J (right image, color represents fiber orientation, right graph is orientation distribution).(c) For three post microtissues, ROIs are made in the center and edge of the tissue as shown (left), each ROI is duplicated (center), and this ROI is used to measure fiber orientation with orientation J (right image, color represents fiber orientation, right graph is orientation distribution). Scalebar 500 µm (left image) and 100 µm (center image).

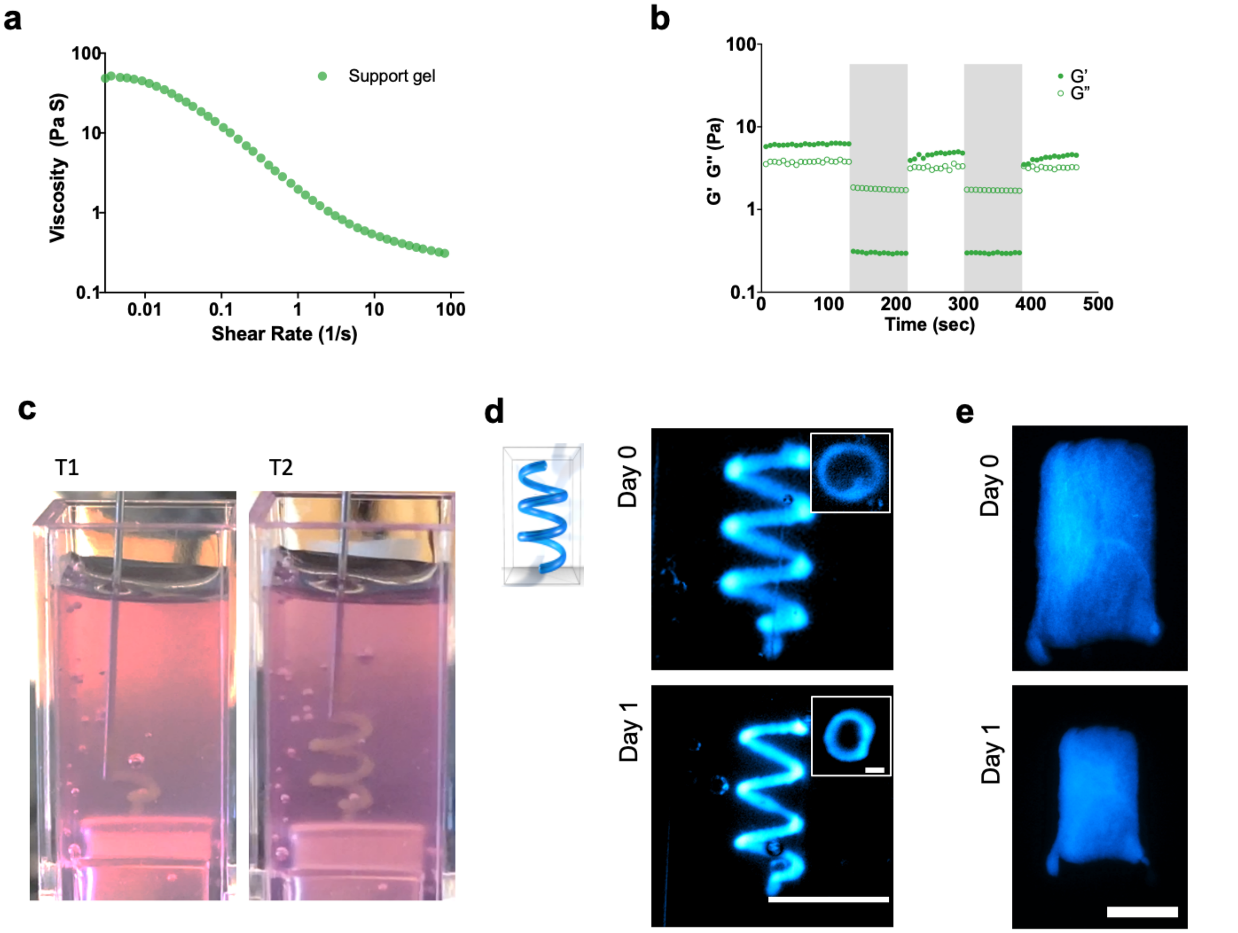


**Figure S14. Rheological properties of support gel for biofabrication studies and 3D printed constructs.** Rheological characterization of support bath hydrogel (0.25 wt % agarose slurry with hyaluronic acid (4 wt%, ~60kDa)) demonstrating (a) shear-thinning properties shown as decreased viscosity with continuously increasing shear rates (0–100 $s^{-1}$) and (b) self-healing properties shown as the recovery of storage modulus during repeated periods of low (1% strain, 1 Hz) and high (shaded, 500% strain, 1 Hz) strain cycles (profiles are representative of n=1 sample). (c) Digital image of printing process in support bath at two time points (T1, T2) (images representative of n=3 biologically separate prints). (d) Representative digital images of spiral shaped prints on day 0 and day 1 after printing, inset shows top view of spiral print at these times (images representative of n=3 biologically independent experiments, each with 3 separate samples, scalebar 5 mm, 1mm for inset). (e) Representative widefield fluorescent images of rectangular prints on day 0 and day 1 after contraction (images representative of n=2 biologically independent experiments, each with 2 separate samples, scalebar 1 mm).

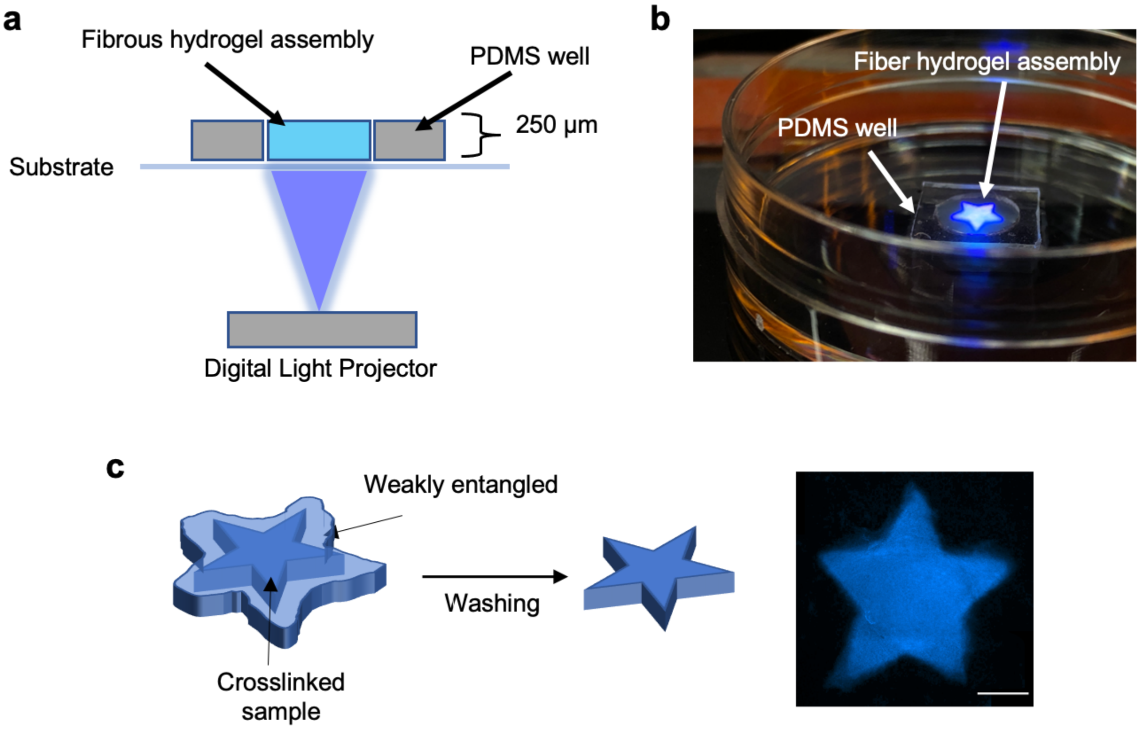


**Figure S15. Photopatterning with digital light projector.** (a) Schematic of photopatterning process and (b) digital image during photopatterning. (c) Post-processing of photopatterned construct to obtain printed shape where a weakly entangled fiber assembly (light blue) around the crosslinked structure (dark blue) is washed to release the photopatterned construct (images representative of n=3 biologically independent experiments each with 2 separate samples, scalebar 1mm).
